# Computational design of structurally diverse antibody repertoires with drug-like properties

**DOI:** 10.64898/2025.12.10.693474

**Authors:** Ariel Tennenhouse, Rebecca Wilen, Sveta Gaiduk, Roman Kamyshinsky, Sarah Borni, Mazal Faraj, Eliane H. Yardeni, Moshe Goldsmith, Inbar Cahana, Shira Albeck, Tamar Unger, Rony Dahan, Ron Diskin, Sarel J. Fleishman

**Affiliations:** Department of Biomolecular Sciences, Weizmann Institute of Science, Rehovot, Israel; Department of Chemical Research Support, Weizmann Institute of Science, Rehovot, Israel; Department of Chemical and Structural Biology, Weizmann Institute of Science, Rehovot, Israel; Department of Systems Immunology, Weizmann Institute of Science, Rehovot, Israel; Protein Analysis Unit, Department of Life Sciences Core Facilities (LSCF), Weizmann Institute of Science, Rehovot, Israel; Structural Proteomics Unit, Department of Life Sciences Core Facilities (LSCF), Weizmann Institute of Science, Rehovot, Israel

## Abstract

Animal immunization is the prevalent strategy for discovering antibody therapeutics^1^, but it is a lengthy and poorly controlled process. As an alternative, synthetic antibody repertoires deliver antibodies without animal welfare concerns, but the resulting antibodies often fail to exhibit “drug-like” biophysical properties^1,2^. Modern repertoires have improved developability by using a handful of frameworks with desirable biophysical properties, but at the cost of reduced structural diversity^3–6^. We developed a principled structure- and energy-based strategy, called CADAbRe, to navigate the complex tradeoffs between developability and structural diversity in repertoire design. The designed repertoire comprises billions of antibodies that are predicted to be stable and foldable, built from hundreds of different frameworks and hundreds of thousands of designed CDR H3s. We also developed an economical and scalable strategy for synthesizing large antibody repertoires, and as a proof of concept, designed and synthesized a 500-million variant phage display repertoire. Selections against four unrelated targets produced structurally diverse binders that exhibited drug-like properties, two of which were readily formatted as bispecifics for functional studies. Furthermore, one of the binders targets a challenging, highly charged surface. The proof-of-concept repertoire is available for academic research. We envision that the CADAbRe approach and repertoire will accelerate and rationalize antibody discovery while addressing animal-welfare concerns^7^.

## Main

Antibody engineering has revolutionized clinical care for a wide range of diseases, with over 100 antibody-based biologics in clinical use and hundreds more undergoing trials^8^. Most antibody therapeutics have been generated using animal immunization^1,9^. However, this is a lengthy and poorly controlled process that depends on a nonhuman animal immune system. For the past three decades, synthetic antibody repertoires have provided an alternative for discovery. These repertoires may be used to discover human-like antibodies that target, in principle, any desired molecule quickly, cheaply, in a controlled *in vitro* setting, and without animal-welfare concerns^10,11^. Increased sophistication in repertoire design has enabled routine discovery of human antibody binders for basic and applied biomedical research^3,12–14^. Yet, despite the widely held expectation that synthetic repertoires would supplant animal immunization^15^, 90% of therapeutics approved in recent years have been sourced from animals^1^. Animal immunization continues to dominate discovery workflows because synthetic antibodies are more often associated with “developability” liabilities, such as low specificity, stability, and serum half-life, than ones from animals^16–18^.

A common strategy to mitigate developability risks is to construct repertoires from a handful of human antibody frameworks that are used in FDA-approved therapeutics^3,4,13^ or exhibit favorable developability properties^6,14^. To compensate for the low framework diversity, the complementarity-determining regions (CDRs), which mediate interactions with the antigen, are typically diversified by recombining CDR sequences from human donors^3^ or introducing random combinations of mutations observed in natural antibodies^5,12^. Modern libraries built using these approaches now deliver developable, high-affinity antibodies^3–6^. However, restricting framework diversity necessarily constrains the space of tolerated CDR sequences and structures, as antibody CDRs feature intricate stabilizing interactions with the framework^19–21^, and diversifying the frameworks and CDR structures can improve the potential to discover binders of challenging epitopes. An additional concern is that many advanced synthetic repertoires are not available for academic research, limiting innovation in biomedicine and the ability to rigorously evaluate the performance of the repertoires^22^. Hence, there is an urgent need for synthetic antibody repertoires that are structurally diverse, developable, and available for academic use.

Recent structure-based methods have successfully optimized antibodies and other biologics, but only on a case-by-case basis^21,23–28^. In addition, *de novo* binder design methodology has made significant advances using deep-learning approaches, including in antibody design^29,30^, but has only been successful in targeting largely hydrophobic surfaces^31^. Moreover, *de novo* antibody design often relies on library screening, requires subsequent affinity maturation, and does not necessarily produce antibodies with “drug-like” properties. By contrast, natural antibodies can recognize a wide range of surfaces^32^, including polar ones^33^, enhancing specificity and the ability to target functional sites. Furthermore, display-based antibody discovery is routinely used by labs around the world to quickly isolate binders against diverse antigens. Thus, combining the ability of structure-based methods to design stable human antibodies with the speed and efficiency of repertoire screening is likely to address pressing needs in antibody discovery workflows.

Here, we present the first structure and energy-based strategy for antibody repertoire design, which we call **C**ombinatorial **A**ssembly and **D**esign of **A**nti**b**ody **Re**pertoires (CADAbRe). CADAbRe selects diverse human antibody germline genes that are predicted to fold stably when combined, accessing antibody gene combinations that have been eschewed in current synthetic repertoires. It then designs sequence diversity within the most important CDR for antigen binding, H3, while ensuring that each design is foldable and energetically stable. Thus, CADAbRe is a principled, energy-based strategy for repertoire generation that does not rely on randomization and greatly expands framework and structural diversity relative to current synthetic repertoires.

Current universal repertoires encode 10^8^-10^11^ unique variants, presenting an unprecedented challenge for structure-based library-design^34–36^. To generate billions of designed antibodies, we developed a scalable computational strategy and a streamlined and cost-effective DNA-assembly strategy. As a proof-of-concept, we synthesized a phage-display repertoire comprising over 500 million human antibodies, orders of magnitude more than structure-based repertoires designed to date^34,36^. As a demonstration, we used phage display to select binders against four different antigens, resulting in structurally diverse and specific binders with developability parameters within the range observed in clinical-stage antibodies. Structural analysis demonstrated that one of the binders targeted a highly charged surface, which would likely defy conventional design methods^31^.

### Structure-based repertoire design

CADAbRe uses structure and energy calculations to identify hundreds of diverse and stable antibody V gene combinations and then designs compatible stable and massively diverse CDR H3 sequences. This strategy departs from current repertoire-generation approaches, which address developability concerns by restricting to a handful of V gene combinations known for high developability and introducing randomized or human-derived CDRs^3–6^.

Because there are no practical methods to synthesize billions of full-length custom-designed antibody variable fragments (Fvs), CADAbRe designs sets of gene fragments that can be freely combined while maintaining stability and structural diversity. It comprises three conceptual steps: first, generating diverse sequence combinations of heavy and light human V and J genes to encode all regions of the Fv except CDR H3; second, using structure-based energy calculations to select those gene combinations that are stable in diverse structural contexts; and third, designing diverse and compatible H3 fragments. We decided to use fully human germline genes in all regions except H3, reasoning that leveraging the natural human CDR diversity would minimize immunogenicity risks. Using fully human germlines has the important additional advantage of retaining so-called GRAB motifs, germline-encoded V gene motifs that have been associated with specific binding to polar antigenic surfaces^33^.

We freely combined the sequences of all human antibody V and J germline genes to generate thousands of human antibody sequences, excluding H3. To evaluate the stability of each germline gene combination in different structural contexts, we grafted H3 sequences from 141 diverse experimentally determined Fv structures^37^ into each germline combination to generate germline-H3 combinations (Fig. 1 Step 1). We ensured that the distribution of H3 lengths matched that seen in therapeutic antibodies (Supp. Fig. 1) and generated an atomistic model for each germline-H3 combination using the antibody stability design algorithm CUMAb^21^. CUMAb modeled each germline-H3 combination on the parental structure from which the H3 was derived (see methods), excluding sequences that differed in length from the parental structure. This procedure yielded over 150,000 models based on the parental Fv structures, in which the H3 retained the sequence observed in the parental structure (Fig. 1, step 2), and the rest of the Fv sequence perfectly matched a combination of human germline genes. One-third of the 150,000 germline-H3 combinations failed this energy-filtering stage, providing examples in which the same germline genes exhibited favorable or unfavorable energy depending on the structural context (Fig. 2A). We verified that modeling different germline combinations on different structures resulted in dramatic differences in structure and electrostatic properties in the antigen-binding surface (Fig. 2B), suggesting that the designed repertoire would encode highly diverse antigen-binding specificities. To further mitigate risks from antibody misfolding, we generated model structures starting from each germline-H3 combination using ABodyBuilder2^38^. Combinations that resulted in ABodyBuilder2 or CUMAb models that deviated substantially from their parental structures were flagged as prone to misfolding (Fig. 2C). Over 27 and 13% of the 150,000 sequences failed these two filters, respectively. We labeled each germline-H3 combination based on its predicted energy and foldability, triaging over half of the 150,000 modeled sequences. Cutoffs and the fraction of sequences passing each filter are available in Supp. Table 1.

**Figure 1:**
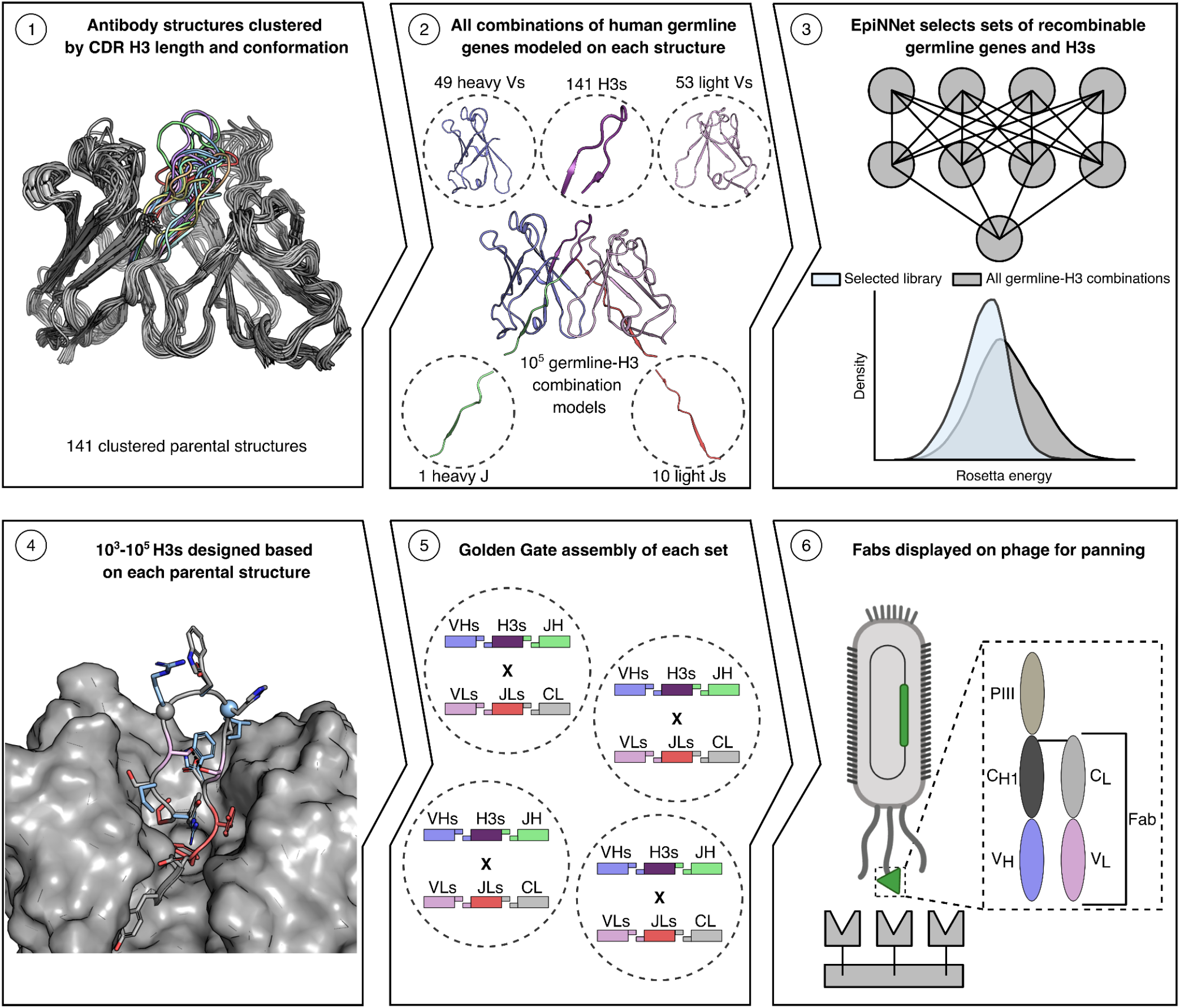
Overview of the CADAbRe repertoire design and synthesis process. High-resolution antibody crystal structures of the variable fragment (Fv) are clustered based on CDR H3 length and conformation (Step 1). All combinations of human germline gene sequences are modeled on each cluster representative and ranked by Rosetta energy (Step 2). The EpiNNet machine-learning approach selects germline genes and parental H3 sequences that can be combined freely to yield variable fragments (Fvs) predicted to be stable and foldable (Step 3). H3 variants are designed for each parental H3 selected in Step 3. Key hydrophobic (red) and hydrogen-bonding (pink) interactions that stabilize the Fv are flagged as immutable. Parental residues are shown in gray, designed residues in blue, and glycines in spheres. The rest of the Fv is shown as a surface (Step 4). Each set of designs is assembled in a one-pot Golden Gate reaction^39,40^ (Step 5). The assembled library is transformed into bacteria and displayed on the surface of phage as antigen-binding fragments (Fabs) for screening (Step 6). Image for Step 6 made with BioRender.

To select sets of germline genes and parental H3 sequences that can be freely combined across diverse structures while maintaining stability and foldability, we applied the EpiNNet machine-learning strategy^34,36^. EpiNNet was previously used to design large repertoires of enzymes^34^, fluorescent proteins^36^, and antibody variants^28^ (Fig. 1 Step 3). It is a one-hidden-layer neural network, which we trained to predict computed stability and foldability metrics based on the component germline genes, parental structures, and the computational results from CUMAb and ABodyBuilder2. The trained network was then used to rank each component, suggesting which are most likely to combine with one another to form stable and foldable full-length Fvs. We used EpiNNet to generate ten sets of combinable fragments that contained the most parental H3 sequences, resulting in 35,000 unique Fv sequences, 85% of which were predicted to be stable and foldable (Fig. 2D and E). The ten sets contained 60 parental structures and over 700 unique light-heavy V gene pairs (Supp. Tables 2 and 3), two orders of magnitude more V gene pairs than recently described synthetic repertoires^3,5^. These 35,000 selected sequences comprised fewer than 25% of the 150,000 modeled sequences and fewer than 4% of the nearly 1 million possible combinations of germline genes and parental structures, significantly focusing on antibodies that are more likely to fold stably into known structures. Despite stability and foldability filtering, the repertoire comprises a much greater diversity of V gene combinations than current synthetic repertoires (Fig. 2F), thus balancing structural diversity, stability, and foldability.

**Figure 2:**
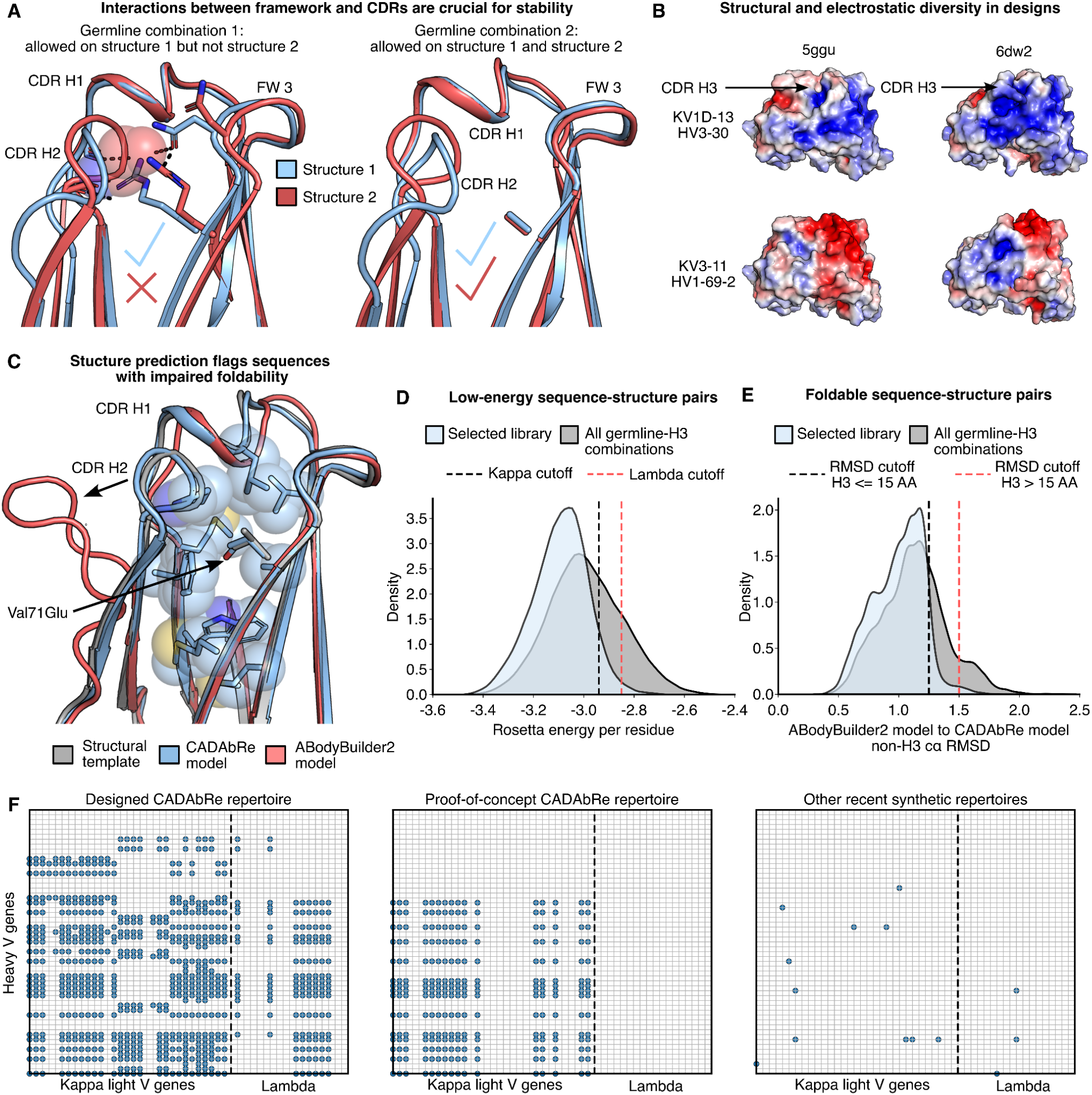
Selection of diverse, stable, and foldable antibody frameworks. (**A**) An example of two germline gene combinations modeled on two different structures. On structure 2, an Arg in germline gene combination 1 forms a steric overlap with backbone atoms of CDR H1, whereas germline gene combination 2 is predicted to be stable. (**B**) Structure and electrostatic diversity are encoded by the combination of germline gene sequences and structures. Electrostatic maps (blue - positive; red - negative) of two germline gene combinations modeled onto two structures within one set. The view shown is looking directly at the CDRs. CDR H3 sequence is identical within each presented structure. (**C**) An example of a germline-H3 combination flagged by ABodyBuilder2^38^ as non-foldable. A Val to Glu mutation creates a buried charge, which is unfavorable. ABodyBuilder2 predicts CDR H2 in a dramatically different conformation, likely to accommodate this buried charge. (**D**) Rosetta energy distributions normalized by protein size. (E) Distributions of the Cɑ distances between the CADAbRe and ABodyBuilder2 models in Fv regions outside of H3. (F) V gene pairs (excluding alleles) present in the complete designed CADAbRe repertoire, the synthesized, proof-of-concept repertoire, and four recent synthetic repertoires^3–6^.

We next used Rosetta atomistic design calculations^41^ to massively diversify each selected H3 based on its parental structure, while maintaining stability and foldability. Because H3 is structurally irregular, diverse, and is the principal contributor to antigen binding, reliable design of H3 is challenging and especially critical for effective repertoires^22^. To support their irregular conformations, H3 structures are stabilized by intricate molecular interactions within themselves and with other parts of the antibody, including the light chain. Maintaining these interactions is key for antibody stability, affinity, and specificity, and sequence randomization, which is commonly used to build conventional repertoires, is very likely to eliminate such stabilizing interactions. We therefore applied the LAffAb CDR atomistic design process^28^ to the parental structures to generate multipoint H3 mutants while retaining the amino acids that make critical contributions to H3 stability (Fig. 1 Step 4, Fig. 3A, Supp. Fig. 2). This procedure automatically identified and held constant positions that make hydrophobic or polar interactions crucial for foldability. It then used Rosetta atomistic calculations to define a sequence space of energetically tolerated mutations, finally ranking combinations of mutations to generate low-energy multipoint mutant designs.

**Figure 3:**
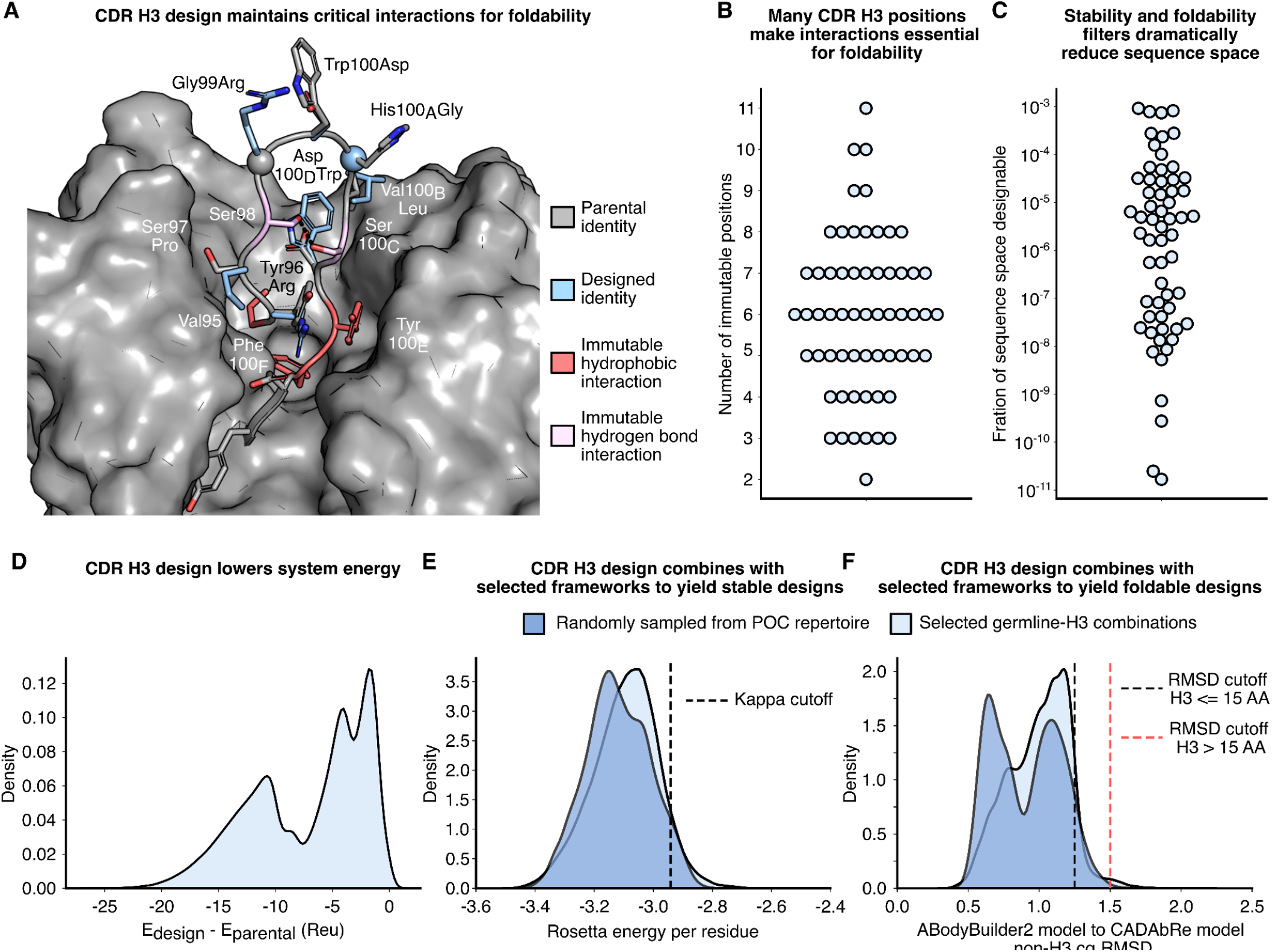
CDR H3 design produces stable and foldable antibodies. (**A**) An example of a specifically designed H3. Ser98 and Ser100_C_ are held fixed due to the stabilizing hydrogen bond (pink). Val95, Try100_E_, Asp101, and Phe100_F_ are fixed due to their interactions with the remainder of the Fv (red). Glycines are shown as spheres. Two additional examples of designed H3s are shown in Supp. Fig. 2. (**B**) Number of positions in each of 60 designed H3s held constant due to essential stabilizing interactions. (**C**) Fraction of the sequence space from per-position residue frequency data that remains after LAffAb filtering for each of 60 designed H3s. (**D**) Rosetta energy distributions of designs compared to the energy of the parental structure for the 19 H3s in the proof-of-concept repertoire. (**E**) Rosetta energy distributions normalized by protein size for 100,000 randomly sampled sequences from the proof-of-concept repertoire compared to the selected germline-H3 combinations. (**F**) Distributions of the Cɑ distances between the CADAbRe and ABodyBuilder2 models in Fv regions outside of H3 in 100,000 randomly sampled sequences from the proof-of-concept repertoire compared to the selected germline-H3 combinations.

The LAffAb filtering of individual mutations for foldability greatly reduced the number of designable positions and the tolerated sequence space of each H3 compared to per-position residue frequency data used in previous repertoire-design efforts^42^ (Fig. 3B and C). In fact, for each H3, the fraction of the tolerated sequence space according to our calculations is at most one thousandth and as little as 2 x 10^-11^ of the space allowed when considering only per-position propensities used to construct some repertoires^12,42^. Despite these restrictions, the space of allowed mutations based on the 60 parental structures comprises millions of sequences. This highlights how structure-based calculations may dramatically focus repertoires on a subspace that is more likely to be stable and foldable while maintaining high sequence diversity. To improve the developability of the designed H3s, methionine and cysteine, which are prone to oxidation, and N-linked glycosylation motifs were not allowed in design. Each H3 exhibited different allowed mutations, reflecting the irregularity and structural diversity of the CDRs (compare Fig. 3A and Supp. Fig. 2).

As a proof of concept, we selected the largest of the ten sets, containing 272 V gene pairs from the IGHV1 and IGHV3 heavy chain and IGKV1, IGKV3, IGKV6 kappa light chain subgroups. We enriched this pool with human germline alleles of heavy and light V genes homologous to those in the set, reasoning that including homologous alleles adds diversity in the CDRs while maintaining the overall features of each germline gene. This resulted in a total of over 450 light-heavy V gene pairs (Supp. Tables 4 and 5), 5 light Js (Supp. Table 6), and 19 diverse parental Fv structures (Supp. Table 8). The allowed H3 mutations for each of these 19 structures were combinatorially enumerated and 240,000 unique and low-energy sequences were selected (Fig. 3D, Supp. Table 9) to favor stability and structural compatibility with other parts of the Fv. In this case, the H3 design was primarily restrained by the number of oligos that could be synthesized, suggesting that the CADAbRe strategy could be scaled to generate much larger repertoires without compromising stability and foldability. The designed H3s were then grafted into the germline combinations generated above, resulting in over 500 million antibody sequences that were designed to be stable and foldable.

We verified that the H3 designs were compatible with all selected germlines by randomly sampling 100,000 designs, modeling each sequence on its parental structure using CUMAb, and, in parallel, generating model structures using ABodyBuilder2 (Fig. 3E and F). Over 90% of the randomly sampled designs passed all CADAbRe filters, confirming that the hierarchical strategy of designing the V gene combinations separately from the H3 diversity generates stable and foldable full-length sequences.

### Scalable assembly of antibody repertoires

To address the challenge of synthesizing a repertoire of this size and complexity, we exploited the fact that diversity is generated through the combination of V and J gene segments (under 100 genes for the entire designed repertoire) and hundreds of thousands of short, designed CDR H3s. This built-in modularity enables rapid, accurate, and cost-effective assembly from oligo pools (for H3) and custom-synthesized V and J genes, similar to reports for other types of proteins^43^ (Fig. 1, Step 5; the assembly process is summarized in Supp. Fig. 3). Furthermore, the fragments can be reused in future assemblies, lowering assembly cost. V and J genes were flanked by suitable sites for Golden Gate cloning^39^ and cloned into plasmids by Twist Bioscience. The light and heavy chains were assembled separately, amplified, and ligated (Supp. Fig. 3A-C). The resulting product, encoding the designed antibodies as IgG1 antigen-binding fragments (Fab), was then ligated into the pComb3x phagemid^44^. We assembled the 5 x 10^8^ designs at a DNA cost of less than $30,000 (under $10^-4^ per antibody variant) in under 10 days, highlighting the efficiency and scalability of the assembly process. A detailed protocol for repertoire synthesis is available^40^.

The library was transformed into TG1 bacterial cells, resulting in over 2 x 10^9^ colony-forming units (CFUs). As an initial check, 24 colonies were subjected to colony PCR and Sanger sequencing (Supp. Fig. 3D). 23 encoded full-length Fabs, and 22 of these exactly matched a designed variant in amino acid sequence, suggesting that >90% of the library matched a designed sequence.

We applied Oxford Nanopore deep sequencing to assess whether the component pieces assembled uniformly. Sequencing of the unselected repertoire generated over 400,000 high-quality mapped reads, allowing us to interrogate the distribution of the H3 sequences and germline genes that comprise the library. All V genes were present in substantial numbers, with less than tenfold difference in the number of reads between the most and least abundant V gene of each chain (Supp. Fig. 4A and B), and the light J genes were represented uniformly (Supp. Fig. 4C). We also found all 464 light and heavy V gene pairs in the library, with no substantial pairing biases (Supp. Fig. 4D). Sequences derived from each of the 19 parental H3s were present, and the distribution of sequences mapped to each H3 resembled the designed distribution for most (Supp. Fig. 4E). Finally, although the read count was less than twice the number of designed H3s, 70% of the designed H3s had at least one mapped read. We concluded that the assembly process is precise and reliable, producing a close-to-uniform distribution of the designed antibodies.

### Diverse and specific binders

To demonstrate that the proof-of-concept repertoire yields binders against diverse antigens, we panned against four target proteins (Fig. 1, Step 6): hen egg white lysozyme (HEWL), the receptor-binding domain (RBD) of the SARS-CoV-2 Spike protein, mouse FMS-related tyrosine kinase 3 (FLT3), and a solubilized version of the Lujo Virus spike protein. These represent different types of targets, from small (HEWL and RBD) to large and glycosylated (FLT3 and Lujo Virus spike). HEWL has attracted intense interest in antibody engineering over decades, allowing us to compare our results to previous biochemical and structural analyses, whereas antibodies with known amino acid sequences that bind to mouse FLT3 and the Lujo Virus spike are unknown to us. FLT3 is particularly interesting as it is a receptor tyrosine kinase that plays a central role in hematopoiesis and dendritic cell biology, regulating the development and maintenance of multiple immune cell populations^45^. Given its importance in immune homeostasis, FLT3 has emerged as an attractive therapeutic target in both oncology and immune modulation^46^.

For each antigen, 18 to 42 colonies from the output of the final panning experiment were assayed for antigen binding using monoclonal phage ELISA against the target antigen, an off-target antigen, and a blocked plate. 52-100% of tested colonies exhibited binding to each target antigen, and none showed nonspecific binding to an off-target antigen or a blank plate (Supp. Table 10). Hits were Sanger sequenced, and at least one designed binder was identified for each target. 12 of the 13 binders exactly matched a designed sequence (Supp. Table 11), and all 13 showed strong on-target binding in monoclonal phage ELISA (Supp. Fig. 5). The 12 binders were selected from fewer than 150 sequenced colonies, a hit rate similar to that seen in applications of other modern repertoires^22^.

The isolated binders were diverse, deriving from 6 light V genes, 3 light J genes, 8 heavy V alleles (4 unique genes), and 6 parental H3s (Supp. Table 11), resulting in dramatic electrostatic and structural diversity in the antigen-binding surfaces (Fig. 4A). We obtained five unique binders against RBD, two against HEWL, and four against FLT3, demonstrating that the repertoire can generate multiple hits for the same antigen. We compared the germline genes used in these 12 binders to those used in four recent repertoire design efforts^3–6^ (Supp. Table 11). Seven of the 12 binders derived from the IGHV1-69 heavy V gene, which is highly represented in clinical antibodies^4^ and was used in two recently described synthetic repertoires^5,6^. Nevertheless, the seven designed binders based on this germline gene derive from four different human alleles, highlighting the importance of allele diversity. The remaining five binders derive from heavy V genes that are absent from all four recently described synthetic repertoires. Similarly, five of the 12 binders derive from light V genes that are absent from these repertoires, demonstrating that CADAbRe productively uses V genes and V gene combinations that state-of-the-art repertoires do not access.

**Figure 4:**
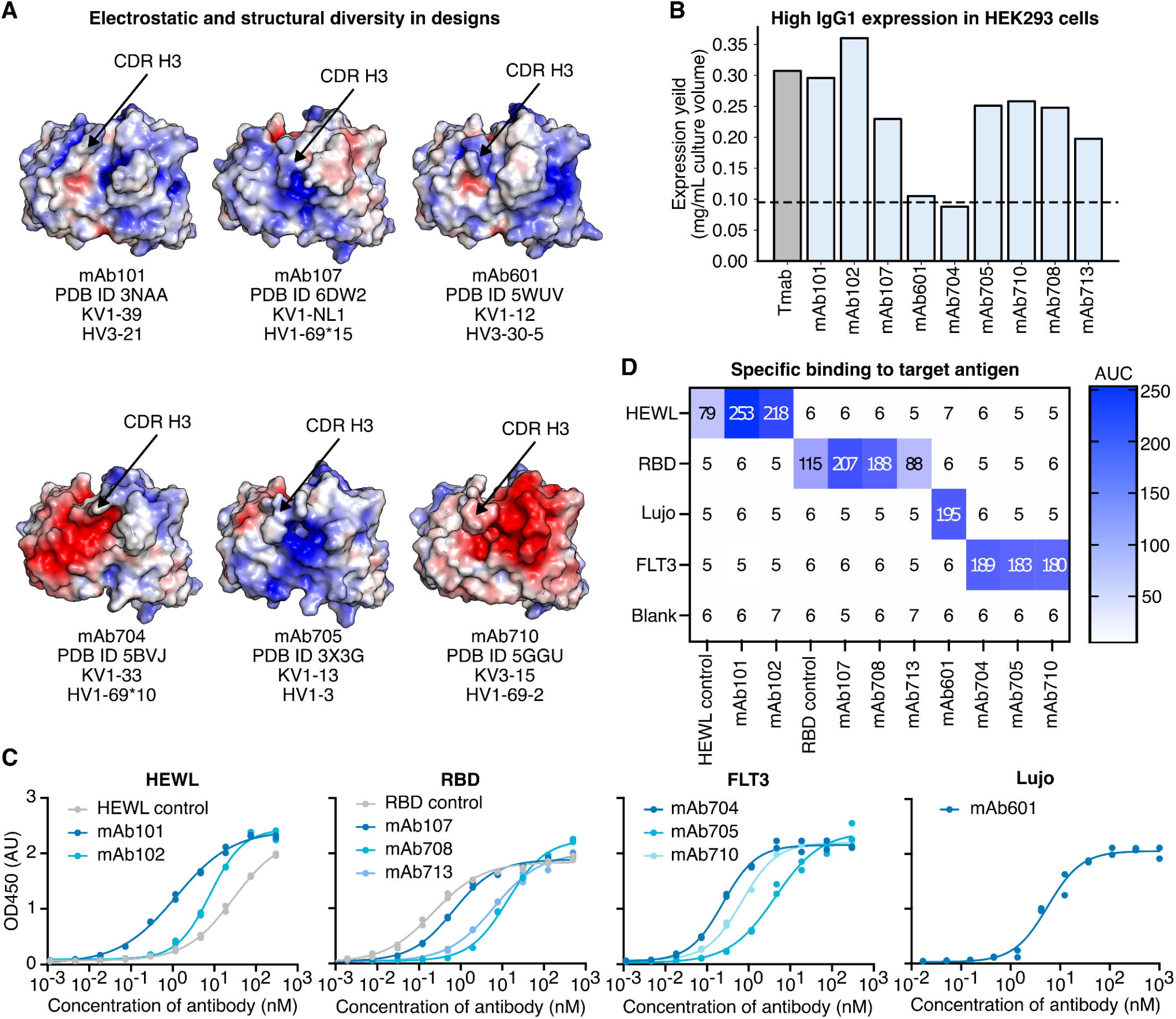
Structural and biochemical analysis of selected designs. (**A**) Selected designs derive from different V gene pairs and parental H3s. Shown are the solvent-accessible surfaces of six selected antibodies colored according to electrostatic potential (blue - positive; red - negative) based on ABodyBuilder2^38^ predicted structures. The view shown is looking directly at the CDRs. (**B**) Expression yields based on a single measurement. Tmab (Trastuzumab) is a control antibody expressed in the same conditions as the designs. The black dashed line represents the historical average of yield in the same experimental conditions (Twist Bioscience). (**C**) ELISAs of the selected designs and controls. Shown is a representative experiment from two replicates. Dots represent the values of two technical replicates, and the lines represent the fit. Fit EC50 values can be found in Supp. Table 12. There are no known antibodies targeting FLT3 and Lujo to serve as positive controls. (**D**) The binding signal was measured using ELISA. Values are the area under the curve of a titration at three concentrations (1, 10, and 100 nM) of each antibody against immobilized antigen. Shown is a representative experiment from two replicates. Curves shown in Supp. Fig. 7. (**C-D**) HEWL control is a previously humanized version of the D44.1 antibody^21^, and RBD control is Cilgavimab.

We expressed nine designs with unique CDR H3s (at least three distinguishing mutations), prioritizing designs with higher monoclonal phage ELISA signal, as full-length human IgG1, and all expressed cleanly (Supp. Fig 6). Eight exhibited yields above the average of constructs expressed in this system, and six exhibited yields more than twice this average (Fig. 4B). The designs exhibited EC50s of 0.3-11 nM (Fig. 4C, Supp. Table 12), and no binding to any of the noncognate antigens (Fig. 4D, Supp. Fig. 7). In two cases for which antibodies were known (RBD and HEWL), the EC50 values of the best designs were within threefold (RBD) or better (HEWL) than the known binder. We concluded that CADAbRe can produce stable and specific binders against diverse targets.

Given the therapeutic interest in FLT3, we reformatted the three FLT3 binders as bispecific antibodies. Two of the three bispecific constructs expressed cleanly (over 85% purity in size-exclusion chromatography) and all three bound FLT3 (Supp. Fig. 8). Bispecific constructs are widely used in modern therapeutic design, but the increased structural complexity of this format is often associated with low developability^47^. Thus, our observations suggest that the repertoire can provide useful starting points for designing complex antibody constructs.

We also measured binding signal by surface-plasmon resonance (SPR) for all hits, either coupling the antibody to the chip when the antigen is a monomer or, in the case of the Lujo spike (a trimer), coupling the antigen and using the antibody Fab as an analyte. All antibodies showed a clear binding signal (Supp. Fig 9, Supp. Table 13). Five measurements could be reliably fit to a 1:1 binding model, with affinities 8 and 13 nM (anti-HEWL), 76 and 238 nM (anti-FLT3), and 691 nM (anti-Lujo), and the other antibodies exhibited fast on and off rates (Supp. Table 13). The affinities, spanning single to triple digit nanomolar dissociation constants, are typical of those obtained from synthetic repertoires and animal immunization^48^. Specifically, HEWL has been a target of benchmarking by other modern synthetic repertoires, which achieved affinities similar to ours (0.5-500 nM)^49^. Finally, we are not aware of antibodies of known sequence that bind to the Lujo spike and FLT3, and the spike protein is hyperglycosylated, presenting a special challenge for binding by antibodies. We conclude that CADAbRe can produce novel antibodies against challenging targets, and that in some cases, these match the affinities of antibodies derived from animal immunization and synthetic repertoires.

### Developable binders

In addition to affinity and specificity, antibodies are ranked according to developability using *in vitro* assays before being nominated as clinical leads. These tests correlate with important therapeutic properties, such as serum half-life, immunogenicity, and aggregation propensity^16,50^. Although developability liabilities are observed even in recently approved antibody therapeutics^16^, molecules that present minimal liabilities are preferred as therapeutic leads. We applied a suite of common developability assays to the nine designs, measuring thermal stability, non-specific binding, self-association, and hydrophobicity (Fig. 5, Supp. Fig 10-17, Supp. Table 14). As a benchmark, we used 12 clinical-stage antibodies that exhibit a range of developability profiles and span different stages of therapeutic development^16^ (Supp. Table 15). For instance, the benchmark contains two FDA-approved antibodies, Adalimumab and Pembrolizumab, which were shown to exhibit zero or several developability liabilities, respectively^16^, providing useful comparisons.

**Figure 5:**
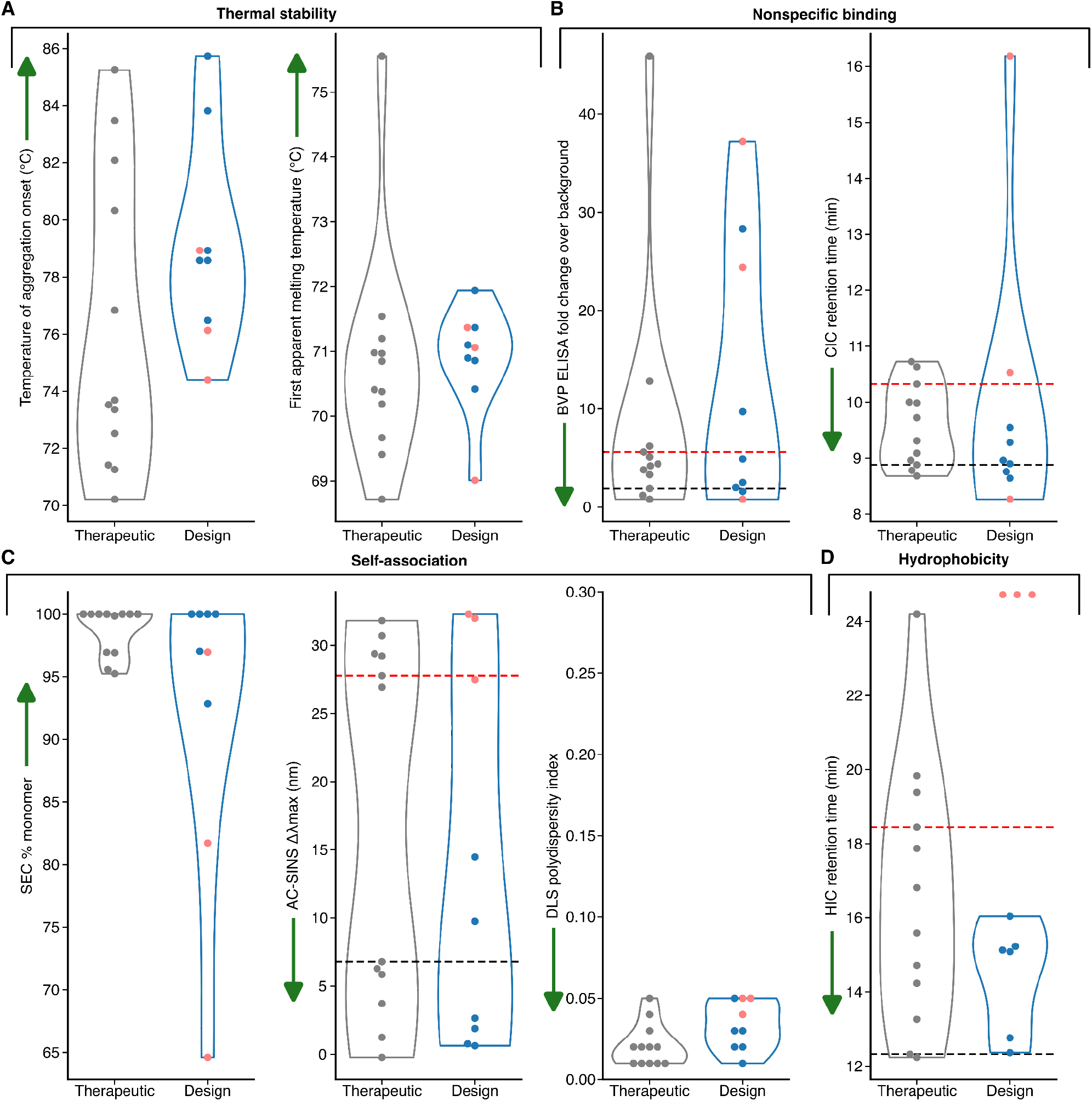
Developability assessment of isolated antibodies compared to clinical-stage controls. Values for each experiment are plotted for the 12 control antibodies and nine isolated designs formatted as IgG1. The designs are colored blue, except for the three designs exhibiting high hydrophobicity (red). Green arrows indicate the desirable direction for each assay. Values for Adalimumab and Pembrolizumab are plotted as black and red dashed lines, serving, where relevant, as lower and upper thresholds, respectively. (**A**) Thermal stability measured by nanoDSF. Dots represent means of two technical repeats. (**B**) Non-specific binding measured using BVP ELISA (dots represent means of four technical repeats) and CIC (based on one measurement). (**C**) Self-association measured using SEC (based on one measurement), AC-SINS (dots represent means of four technical repeats), and DLS (dots represent means of two technical repeats). (**D**) Antibody hydrophobicity measured using HIC (based on one measurement). The three designs exhibiting high hydrophobicity did not elute from the column and are placed at the y-axis maximum. nanoDSF: nano differential scanning fluorimetry; BVP: baculovirus particle^52^; CIC: cross-interaction chromatography^53^; SEC: size-exclusion chromatography; AC-SINS: affinity-capture self-interaction nanoparticle spectroscopy ^54–56^; DLS: dynamic light scattering; HIC: hydrophobic interaction chromatography.

All designs and controls exhibited high apparent thermal stability in aggregation (*T_agg_* > 70°C) and unfolding experiments (*T_m_* > 68°C) (Fig. 5A, Supp. Fig. 10 and 11). Seven designs exhibited over 90% purity by SEC (Fig. 5C, Supp. Fig. 14), suggesting that most designs have stability and purity similar to therapeutic candidates (all controls exhibited over 95% purity). Most designs exhibited low nonspecificity (5 and 7, respectively in BVP and ClC; Fig. 5B, Supp. Fig. 12 and 13); high values in these parameters are predictive of fast *in vivo* clearance^51^ and therefore more accurately reflect specificity than ELISA experiments against a fixed panel of proteins (Fig. 4D). Six also exhibited low self-association (AC-SINS; Fig. 5C, Supp. Fig 15, Supp. Table 16) and hydrophobicity (HIC; Fig. 5D, Supp. Fig. 16), and all antibodies exhibited low polydispersity (DLS) (Fig. 5C, Supp. Fig. 17). Thus, most designs exhibited few or no liabilities with a few flagging for hydrophobicity and nonspecific binding.

We visually inspected the structural models of the three antibodies that were flagged for hydrophobicity, noting that they comprised a Phe or Trp residue at the solvent-exposed tip of CDR H3, in contrast to the other isolated designs (Supp. Fig. 18). Such aromatic residues may, in some cases, be necessary for high-affinity binding. In fact, two of the flagged antibodies targeted the RBD, and other studies reported high hydrophobicity in anti-RBD antibodies^3^, suggesting that effective binding to this antigen requires at least some hydrophobicity. An important advantage of CADAbRe is that it is straightforward to constrain the repertoire composition based on empirical evidence. Thus, repertoires currently being designed in our lab avoid exposed aromatics, and future repertoires will incorporate lessons based on new findings, increasing the developability and usefulness of the CADAbRe repertoires.

We concluded that the developability of most designs were on par with clinical-stage antibodies. These results compare favorably with therapeutic antibodies previously isolated from phage-display libraries, which revealed a greater tendency to show developability liabilities than antibodies derived from animals^2,23^. We note that comparing designs to clinical-stage antibodies sets a high bar, as antibodies undergo stringent testing and engineering before being promoted to human trials, and Pembrolizumab (Keytruda) is a blockbuster drug despite its liabilities. Thus, most designs could be viewed as exhibiting drug-like properties, despite having been isolated from a standard panning procedure without undergoing any developability filtering or engineering. Future mining of antibodies from this repertoire will help characterize the overall developability of the repertoire and identify additional areas for improvement of the design methodology.

### Binding to a charged epitope

Protein-protein interfaces are often dominated by hydrophobic interactions, and highly polar surfaces present a challenge for binder discovery and design^31^. By contrast, the germline V gene repertoire encodes so-called GRAB motifs that can bind specifically to polar and charged amino acid sidechains^33^, increasing the range of molecular surfaces that can be targeted. We reviewed the isolated binders for such motifs, finding that the anti-HEWL mAb102 comprises IGHV3-21, which harbors a binding motif for Asp or Glu on CDR H2.

To demonstrate that the CADAbRe repertoire may enable binding to charged surfaces, we determined the cryo-electron microscopy (cryo-EM) structure of mAb102 in complex with HEWL to a resolution of 3.1 Å (Supp. Fig. 19). Despite 57 mutations relative to the parental structure that served as a template for design (PDB entry: 3NAA), including four mutations in CDR H3, the two structures are highly similar (1.2 Å Cɑ root-mean-square-deviation; RMSD) (Fig. 6A center), verifying the accuracy and foldability of the designed antibodies. We noticed that the tips of CDRs H2 and H3 swiveled to form closer interactions with the antigen relative to their position in the parental structure (Supp. Fig. 20), demonstrating that the designed mutations ensure foldability while allowing the CDRs to adapt to the target surface. Furthermore, we found that the designed mutations in H3 stabilize the bound conformation by interacting with other CDRs or relieving clashes with the antigen (Supp. Fig. 20A).

**Figure 6:**
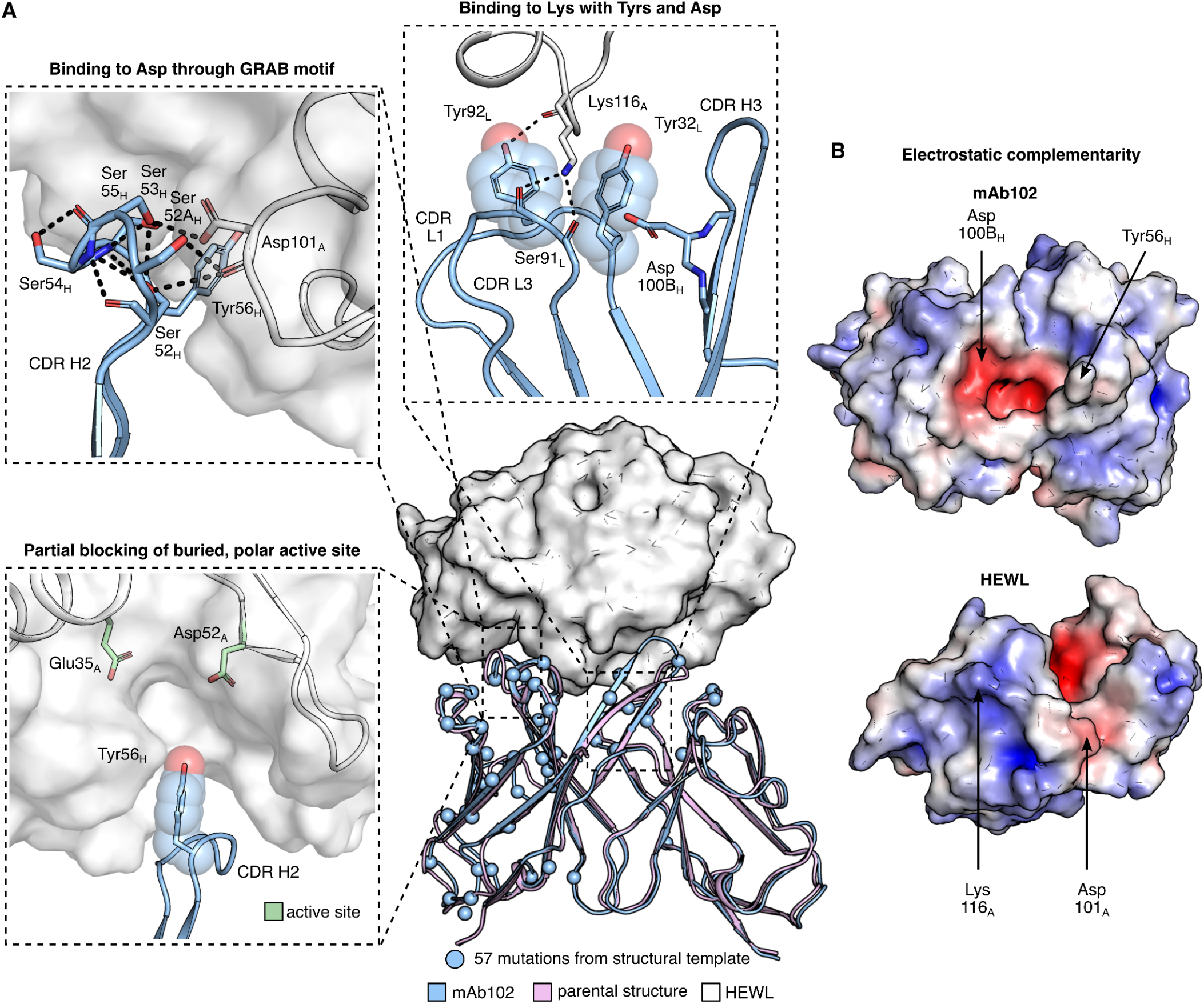
Cryo-EM structure of the mAb102/HEWL complex reveals a highly polar interface. (**A**) The cryo-EM structure recapitulates the parental structure (PDB ID 3NAA) despite 57 mutations (central panel). CDR H2 binds to an Asp on HEWL through a GRAB motif (top-left panel). CDRs L1, L3, and H3 bind to a Lys on HEWL through a pair of Tyrs and an Asp (top panel). Binding to this polar surface enables mAb102 to partially occlude the active site of HEWL (bottom-left panel). HEWL is shown as a surface map in the center and left panels. (**B**) Open book view of the mAb102/HEWL interface, with HEWL rotated by 180° to display the interface. (**A and B**) Subscript L, H, and A denote light chain, heavy chain, and antigen (HEWL), respectively. Antibody position numbering according to Kabat.

Strikingly, the structure reveals that mAb102 binds to two distant charges on HEWL: the known CDR H2 GRAB motif interacts with HEWL Asp101 (Fig. 6A top left), and HEWL Lys116 interacts with two Tyrs (on L1 and L3) and an Asp on H3 (Fig. 6A top and 6B). Likely due to the difficulty of binding two charged residues, this binding orientation is not seen in any other HEWL-binding antibody, to the best of our knowledge. Because charge-mediated binding is critical for specificity, particularly for targeting functional sites, we asked whether the Lys-targeting binding mode is seen in other antibody-antigen complexes. We searched for bound antibody structures in the PDB that comprise similar light chain sequences to mAb102 and exhibit two Tyr residues aligned to those in mAb102 that interact with a Lys sidechain on the antigen. This search yielded six human antibodies that bind to unrelated antigens in which the structure closely recapitulated the interaction with Lys seen in mAb102, comprising Tyr, Tyr, and Asp/Glu (L1, L3, and H3, respectively) (Supp. Fig. 21). This structural analysis extends the notion of GRAB motifs to include complex constellations of amino acids from different germline genes.

Critically, by interacting with two exposed and charged amino acid side chains, mAb102 partially occludes the active site of HEWL (Fig. 6A, bottom left). This demonstrates that by encoding many germline gene combinations, CADAbRe can target complex functional surfaces that would likely defy current synthetic repertoires and *de novo* binder design methodology^31^. Moreover, targeting polar antigenic surfaces reduces the number of exposed hydrophobic residues on the CDRs, contributing to developability. These observations suggest that, in addition to generating universal repertoires, CADAbRe could be customized for interacting with specific challenging surfaces. For instance, repertoires built using V genes that encode GRAB motifs could be used to select binders for polar surfaces, and ones that encode glycan-binding motifs could be used to target glycoproteins^57^.

## Discussion

Synthetic repertoires are a versatile and powerful platform for generating human antibody binders quickly, cheaply, and in experimentally controlled *in vitro* settings. But generating many therapeutic leads demands encoding diverse structures and sequences of highly developable antibodies. Early synthetic repertoires recombined many different V genes to emphasize diversity^58,59^, but these repertoires tended to deliver antibodies that lacked critical developability properties. To address these deficiencies, modern repertoires have been designed with a restricted set of V gene combinations verified for developability. Furthermore, modern repertoires typically undergo *post hoc* experimental filtering to increase developability but at the cost of further restricting diversity. Despite the important advances in generating repertoires with developable antibodies, however, animal immunization, with its associated ethical concerns, has continued to dominate therapeutic lead discovery, with over 1 million animals estimated to be sacrificed annually in the E.U. alone^60^. The continued dominance of immunization is due to the higher diversity, affinity, and developability of antibodies derived from animal immunization, and because access to the best synthetic repertoires remains restricted both in academia and industry^1^. In addition, it is estimated that hundreds of millions of dollars are wasted annually in biomedical research on suboptimal antibody reagents^61^. Addressing the diversity and developability limitations of synthetic repertoires and making them accessible to researchers can therefore contribute significantly to the ethics and productivity of future basic and applied biomedical research.

Despite decades of research on the structural underpinnings^62^ of antibody stability^19^ and activity^63^, CADAbRe is the first repertoire-design approach guided by principles rooted in energy and structure. It navigates the complex landscape of repertoire diversity and developability by restricting to known antibody structures rather than to specific V gene combinations that are considered developable. The current proof-of-concept repertoire is built on 19 diverse template structures, and future repertoires will employ many more structures to increase epitope-targeting diversity. Furthermore, by encoding fully human V genes, the repertoire reduces immunogenicity risks and exploits the innate ability of the human germline to bind specifically to polar and charged surfaces that are likely to defy conventional computational and experimental techniques.

It is encouraging that a design procedure that optimizes only foldability and native-state energy delivered multiple antibodies with developability properties akin to clinical-stage ones against diverse targets. Moreover, the cause of hydrophobicity in some antibodies was readily explained by modeling. Additional screening against different targets will shed light on other areas for improving the repertoire, and the programmability of the repertoire provides a path towards continuous improvement through design-test-learn cycles.

The proof-of-concept repertoire we generated and analyzed verifies our hypothesis that greater V gene diversity can be encoded in synthetic repertoires without sacrificing developability. The affinities we obtained were comparable or lower relative to those obtained from established repertoires, likely due to the small size of the repertoire, which will be substantially enlarged in future iterations. We note that the repertoire produces stable and developable binders with no mutations outside of H3, thus providing excellent leads for affinity maturation processes. The vast space of low-energy human antibody structures and sequences that CADAbRe can design and the streamlined repertoire synthesis approach enable generating larger repertoires that will access even more diversity and yield higher-affinity binders directly from selections. Indeed, the current proof-of-concept repertoire comprises only a fraction of the repertoire we designed. Furthermore, DNA synthesis technologies are rapidly improving, and CADAbRe can flexibly adapt to such advances to encode even larger repertoires.

CADAbRe combines the power of phage display to deliver therapeutic leads quickly and economically with the ability of modern modeling and design approaches to design diverse, stable and foldable antibodies. Its goal is to generate synthetic repertoires that represent a large diversity of human germline genes while producing antibodies that exhibit properties seen in therapeutic leads. We envision that this effort will ultimately produce antibodies with developability features that closely resemble or even exceed those from animal immunization without the associated time, cost, ethical concerns, and requirement to engineer humanized versions. The repertoire is accessible to academic researchers to accelerate the discovery of promising therapeutic leads and research reagents while reducing animal-welfare concerns in biomedical research.

## Methods

### Antibody numbering

Antibody numbering was performed using the AbNum^64^ web server (http://www.bioinf.org.uk/abs/abnum/). Kabat numbering^65^ is used throughout the work.

### Computational methods

All Rosetta design simulations used git v.d9d4d5dd3fd516db1ad41b302d147ca0ccd78abd of the Rosetta biomolecular modeling software, which is freely available to academics at http://www.rosettacommons.org. All RosettaScripts^66^ XMLs can be found at https://github.com/Fleishman-Lab/CADAbRe_public.

#### A database of human antibody germline sequences

Human antibody germline amino acid sequences were retrieved from the IMGT reference database^67^ (downloaded April 19, 2023). Heavy V genes were trimmed to the conserved cysteine before CDR H3 (position 92). To be included in the database, a gene must have a unique amino acid sequence, be annotated as functional (i.e., not partial or reverse-complementary), and not contain N-glycosylation sites (NX[S/T], where X is not Pro). Moreover, a V gene must contain exactly two cysteines, and a J gene must contain zero cysteines. D genes were not considered, and only one heavy J gene was used (IGHJ1*01). IGHJ1*01 was trimmed to the conserved tryptophan after CDR H3 (position 103). Kappa light V genes were considered only if they ended in proline so that the proline could be used to define an overhang for Golden Gate cloning to ligate light V to light J. Generally, the first allele for each gene was included. If the first allele did not satisfy any of the above criteria, a different allele was taken if one could be found that satisfied all the above criteria. This procedure resulted in 49 heavy-chain V-gene sequences, 31 kappa light-chain V-gene sequences, 5 kappa light-chain J-gene sequences, 28 lambda light-chain V-gene sequences, and 5 lambda light-chain J-gene sequences. We combined these genes all-against-all (V and J for both light and heavy chains) for kappa and lambda light chains separately, resulting in 7,595 sequences for kappa light chains and 6,860 sequences for lambda light chains.

#### A database of antibody variable fragment (Fv) structures

Antibody structures were downloaded from the Structural Antibody Database (SabDab)^37^ on September 7, 2020. The “search for a non-redundant set of antibodies” feature on the SabDab web server was used to select a set of paired VH/VL structures with a maximum 90% sequence identity threshold and crystallographic resolution <= 2.8Å. The remaining structures were filtered for those of murine or human origin that were experimentally solved using X-ray crystallography. This procedure resulted in 960 high-quality human or murine antibody structures.

#### Clustering Fv structures by CDR H3 conformation

Variable fragments were extracted from each structure by aligning each structure to the variable fragment from PDB ID 2BRR^68^, which served as a reference. AbDesign^69^ was used to segment the CDR H3 (positions 93-103, 103 included to facilitate structural alignment) of each structure based on the alignment with 2BRR. H3 structures with missing density were purged. The remaining H3 structures were grouped by H3 length (positions 93-102). Only lengths 9-20 and 23 were considered. This matches the lengths seen in clinical-stage antibody therapeutics^70^. This procedure resulted in 259 cluster representatives.

#### Additional filtering of selected Fv structures

The Fv structures of each CDR H3 cluster representative were loaded into PyMOL and visually inspected. Structures with missing density anywhere in the variable fragment were discarded. Additionally, structures that were deposited in the PDB before the year 2000 were discarded. Finally, structures that did not have at least one human light and heavy V gene with matching CDR lengths of non-H3 CDRs were discarded. This procedure resulted in 141 Fv structures.

#### Computational generation of full Fv sequences

For each selected Fv structure, the CDR H3 sequence (positions 93-102) from the parental structure was grafted into all germline combinations with matching light chain type and non-H3 CDR lengths. Sequences containing N-glycosylation sites (NX[S/T], where X is not Pro) were discarded. This procedure resulted in 153,775 unique Fv sequences.

#### Energy-ranking of human germlines across selected structures

The structure of each selected parental antibody Fv was relaxed through cycles of sidechain and harmonically restrained backbone minimization and combinatorial sidechain packing in the entire Fv (see xmls/Relax.xml). Each generated Fv sequence was threaded onto each relaxed Fv structure and relaxed in the same manner with fewer cycles (see xmls/CADAbRe.xml). Each unique Fv sequence modeled on a structure (germline-H3 combination model) was ranked according to the all-atom energy function REF15 ^41^ divided by the number of residues (“energy per residue”) to account for differences in sequence length.

#### Labeling Fv germline-H3 combination model as stable and foldable

Each germline-H3 combination model was assessed computationally for stability by comparing the Rosetta energy per residue to that of each original sequence from each pdb structure modeled onto its own structure using the same procedure. Germline-H3 combination models were labeled as stable if their energy per residue was less than or equal to the mean of the original sequences plus two standard deviations. The foldability of each germline-H3 combination model was evaluated using two strategies. First, the carbonyl C+O RMSD of each CDR of each germline-H3 combination model to the original, relaxed structure was calculated using PyMOL. Germline-H3 combination models were labeled as unfoldable if they had CDR L1 RMSD > 1Å, L2 RMSD > 1.2Å, L3 RMSD > 1.2Å, H1 RMSD > 1.3Å, H2 RMSD > 1.2Å, or H3 RMSD > 1.3Å. These thresholds were determined by visual inspection. ABodyBuilder2^38^ was also used to predict the structure of each sequence, and the Cα RMSD of the predicted structure to the model generated by threading onto the original, relaxed structure was calculated using PyMOL. Sequences that had a non-CDR H3 RMSD > 1.25Å (CDR H3 length <=15) or > 1.5A (CDRH3 length > 15) were labeled as unfoldable. Sequences passing both RMSD thresholds were labeled as foldable.

#### Selecting a set of recombinable human germline genes and structures

Germline-H3 combinations were divided into sets based on their non-H3 CDR lengths. For the ten sets containing the most structures, EpiNNet^34,36^ was applied to select for pools of germline genes and structures that recombine all vs. all to give a large fraction of germline-H3 combinations labeled as stable and foldable. For most pools, this resulted in >80% of the germline-H3 combinations labeled as stable and foldable.

#### CDR H3 design of selected structures

For each selected structure, positions 95-102 were allowed to be designed using the LAffAb CDR design process^28^. Positions 93 and 94 were held constant as position 93 likely makes limited contributions to binding, and 94 is well-known to make stabilizing interactions within H3^71^. Allowed mutations were initially identified based on per-position residue frequency data^42^. For each H3 category (“short”, “medium”, and “long” defined by Prassler et al^12^.), identities were allowed at each position if they had a frequency >= 4%. Positions labeled in Prassler et al. as “E” and “F” (“medium” CDRH3 length) or “N” and “O” (“long” CDRH3 length) were considered to be the last two residues before positions 101 and 102. Position 101 was maintained as Asp if it was Asp in the original structure, and position 102 was allowed to be His, Leu, Val, Ile, Tyr, Ser, or Pro (the identities found in position 102 in human heavy J germline genes). Methionine and cysteine were not introduced in any position to avoid methionine oxidation and unwanted disulfide bonds, respectively. N-linked glycosylation sites (NX[S/T], where X is not Pro) were not allowed.

Positions 95-102 were each mutated to Ala and scored in Rosetta using the REF15 score function^41^ and a version of the REF15 score function that only evaluates hydrogen bonds (see xmls/alascan.xml and xmls/alascan_Hbond.xml). Positions with +3.5 REU in REF15 energy or +0.6 REU in REF15 hydrogen bond energy are held constant. Each identity allowed by the per-position frequency data in all other positions is then individually modeled and scored (see xmls/filterscan.xml). All mutations that have <= +4 REU are then enumerated combinatorially for each structure (see xmls/H3_combinatorial_enumeration.xml). If this results in a number of combinations too large to be enumerated, the threshold is reduced to +3 REU. Designs with Rosetta energy less than or equal to the Rosetta energy of the original sequence for each structure are considered.

#### Clustering of selected CDR H3 designs

The CDR H3s were ordered as a cloned oligo pool from Twist, consisting of 240,000 oligos. This section deals with how the 240,000 oligos were allocated to H3 designs from different parental structures. Parental structures with more than 100,000 designs with Rosetta energy less than or equal to the original sequence were clustered by two mutations. Parental structures with fewer than 100,000 designs with Rosetta energy less than or equal to the original sequence were not clustered. To account for some parental structures being easier to design than others, the following log-based normalization strategy was adopted.

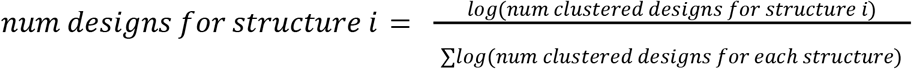

Structures with less than 10,000 LAffAb-generated designs were not included in this log correction, and all designs with energy <= that of the original sequence for these structures were included.

#### DNA sequences of designed CDR H3s

Each designed CDR H3 encompassed positions 93-102, with 93 and 94 maintained as in the parental structure. These amino acid sequences were reverse-translated using the back_translate method in CodonHarmony. The resulting DNA sequences were then codon optimized for expression in *E. coli* using DNA Chisel^72^. DNA chisel was constrained to avoid *SapI, BsaI, BbsI, and EcoRV* sites, which are used in library assembly. Additionally, DNA Chisel was constrained to avoid the following motifs: [GC][GC][GC][GC][GC], TTTTTT, AAAAAA, and to ensure the GC fraction of the resulting sequence was between 0.4 and 0.6. The random seed in NumPy^73^ was set to 123, so the sequences will be reproduced upon reruns. The codon “TGC” (encoding for cysteine) was appended to the 5’ end of each sequence, and the sequence “TGGGGTCAAGGCACG” (encoding for WGQGT) was appended to the 3’ end of each sequence. “TGC” and “ACG” are gates that will be used in the Golden Gate^39^ assembly. The resulting sequence encodes for positions 92-107.

A *SapI* site is added to the 5’ end (“GCTCTTCC”) and 3’ end (“CGAAGAGC”) of each sequence. Then, an *EcoRV* site (“GATATC”) is added to the 5’ and 3’ ends of the sequence. A six-base-pair padding sequence (“GACATT”) is added between the 5’ *EcoRV* and *SapI* sites. A longer padding sequence is added between the 3’ *SapI* and *EcoRV* sites. This padding sequence consists of a unique primer for each parental structure, selected from a set of primers experimentally validated to be orthogonal to each other^74^. If the resulting sequence is longer than 120 base pairs, the primer is shortened until the total sequence is 120 base pairs. If the resulting sequence is shorter than 120 base pairs, additional base pairs are added until the total sequence is 120 base pairs. Each final sequence is thus constructed as follows:

*EcoRV* - pad - *SapI* - TGC (Cys) - CDR H3 coding sequence - TGGGGTCAAGGC (WGQG) - ACG (Thr) - *SapI* - primer - *EcoRV*

The 240,000 sequences were then ordered as an oligo pool cloned into the pTwist Kan High Copy v2 plasmid by Twist Bioscience. Cloned oligo pools were used because they alleviate the need for double-stranding reactions, which have a high error rate, and because cloned oligo pools are straightforward to amplify by bacterial transformation. In principle, single-stranded oligos could also be used, which would decrease the cost of the library, likely at the cost of decreased fidelity.

#### Selection of V gene alleles to add or remove

Enumerating all combinations of V gene alleles across all germlines would be computationally expensive, so only one allele per V gene was enumerated. After the V genes were selected, the V gene alleles of each V gene were visually inspected. Alleles encoding additional diversity in the CDRs that did not interact with CDR H3 or the framework were added to the set to enrich diversity without impacting interactions with other parts of the antibody. Additionally, CDR1 and 2 were concatenated for each V gene. V genes that did not have a unique CDR1+2 were removed.

#### DNA sequences of light V germline genes

The C-terminal Pro in each light V gene was trimmed. Each light V sequence was then reverse-translated using the back_translate method in CodonHarmony. The resulting DNA sequences were then codon optimized for expression in *E. coli* using DNA Chisel^72^. DNA chisel was constrained to avoid *SapI, BsaI, BbsI,* and *EcoRV* sites, which are used in library assembly. Additionally, DNA Chisel was constrained to avoid the following motifs: [GC][GC][GC][GC][GC], TTTTT, AAAAA, and to ensure the GC fraction of each 10-base-pair window was between 0.3 and 0.7. DNA Chisel was also constrained to avoid 100% matches to any 15-base-pair window in any previously codon-optimized light V gene. The random seed in NumPy^73^ was set to 123, so the sequences will be reproduced upon reruns. The DNA sequence encoding for the amino acids RFSGSG was replaced with CGATTTTCTGGCTCTGGCA so that each light V gene contains an internal primer binding site. Another primer binding site (“CCAGCCTGATAAGTAGCACCGAAGACCCTGCC”), a *BbsI* site (“GAAGACCC”), and the gate to eventually ligate into the modified pComb3X^44^ vector (“TGCC”) were added to the 5’ end of each light V sequence. Then, a gate to ligate into the light J plasmids (“GTT”), a *SapI* site (“GCTCTTCC”), and an *EcoRV* site (“GATATC”) were added to the 5’ end of each light V sequence. Finally, a second gate to ligate into the light J plasmids (“CCT”, encoding for proline) and a *SapI* site (“CGAAGAGC”) were added to the 3’ end of each light V sequence. Each final light V sequence was thus constructed as follows:

*SapI* - GTT - *EcoRV* - primer - *BbsI* - TGCC - light V coding sequence - CCT (Pro) - *SapI*

The resulting sequences were ordered as genes cloned into the pTwist Chlor High Copy plasmid by Twist Bioscience.

#### DNA sequences of heavy V germline genes

The C-terminal Cys in each heavy V gene was trimmed. Each heavy V sequence was then reverse-translated using the back_translate method in CodonHarmony. The resulting DNA sequences were then codon optimized for expression in *E. coli* using DNA Chisel^72^. DNA chisel was constrained to avoid *SapI*, *BsaI*, *BbsI*, and *EcoRV* sites, which are all used in the library assembly pipeline. Additionally, DNA Chisel was constrained to avoid the following motifs: [GC][GC][GC][GC][GC], TTTTT, AAAAA, and to ensure the GC fraction of each 10-base-pair window was between 0.3 and 0.7. DNA Chisel was also constrained to avoid 100% matches to any 15-base-pair window in any previously codon-optimized heavy V gene. The random seed in NumPy^73^ was set to 123, so the sequences will be reproduced upon reruns. An *EcoRV* site (“GATATC”) and most of the PelB leader sequence (excluding the 5’ “AT”, “GAAATACCTATTGCCTACGGCAGCCGCTGGATTGTTATTACTCGCTGCCCAACCAGCCATGGCC”) were added to the 5’ end of each heavy V sequence. A constant primer was added when possible to enable PCR for deep sequencing of CDR H3. A gate to ligate to the CDRH3 oligos (“TGC”, encoding for cysteine), a *SapI* site (“CGAAGAGC”), a six-base-pair padding sequence (“GACATT”), and an *EcoRV* site were added to the 3’ end of each heavy V sequence. Each final heavy V sequence was thus constructed as follows:

*EcoRV* - PelB leader - heavy V coding sequence - TGC (Cys) - *SapI* - pad - *EcoRV*

The resulting sequences were ordered as genes cloned into the pTwist Kan High Copy v2 plasmid by Twist Bioscience.

#### DNA sequence of the heavy J germline gene

The part of the IGHJ1*01 amino acid sequence not included in the CDR H3 oligos (LVTVSS) was reverse-translated using the back_translate method in CodonHarmony. The resulting DNA sequence was then codon optimized for expression in E. coli using DNA Chisel^72^. DNA chisel was constrained to avoid *SapI*, *BsaI*, *BbsI*, and *EcoRV* sites, which are all used in the library assembly pipeline. Additionally, DNA Chisel was constrained to avoid the following motifs: [GC][GC][GC][GC][GC], TTTTT, AAAAA, and to ensure the GC fraction of each 10-base-pair window was between 0.3 and 0.7. The random seed in NumPy(Harris et al. 2020) was set to 123, so the sequence will be reproduced upon reruns. An *EcoRV* site (“GATATC”), a six-base-pair padding sequence (“GACATT”), a *SapI* site (“GCTCTTCC”), and a gate for ligating to the CDRH3 oligos (“ACG”, encoding for threonine) were added to the 5’ end of the heavy J sequence. A gate for ligating into the modified pComb3X^44^ vector (“GCTA”) and a *BbsI* site (“CCGTCTTC”) were added to the 3’ end of the heavy J sequence. Next, 300 base-pairs of random DNA were generated by DNA Chisel using the random_dna_sequence function with gc_share=0.45 and seed=2. This 300-base pair segment was optimized by DNA Chisel to avoid *SapI*, *BsaI*, *BbsI*, and *EcoRV* sites, which are all used in the library assembly pipeline. Additionally, DNA Chisel was constrained to avoid the following motifs: [GC][GC][GC][GC][GC], TTTTT, AAAAA, and to ensure the GC fraction of each 10-base-pair window was between 0.3 and 0.7. The resulting 300-base-pair segment of DNA and an *EcoRV* site were added to the 3’ end of the heavy J sequence. The final heavy J sequence was thus constructed as follows:

*EcoRV* - pad - *SapI* - ACG (Thr) - heavy J coding sequence - GCTA - *BbsI* - 300 base pairs of random DNA - *EcoRV*

The heavy J sequence was ordered as a gene cloned into the pTwist Kan High Copy v2 plasmid from Twist Bioscience.

#### DNA sequences of light J germline genes

Each light J sequence was reverse-translated using the back_translate method in CodonHarmony. The resulting DNA sequences were then codon optimized for expression in E. coli using DNA Chisel^72^. DNA chisel was constrained to avoid *SapI*, *BsaI*, *BbsI*, and *EcoRV* sites, which are all used in the library assembly pipeline. Additionally, DNA Chisel was constrained to avoid the following motifs: [GC][GC][GC][GC][GC], TTTTT, AAAAA, and to ensure the GC fraction of each 10-base-pair window was between 0.3 and 0.7. DNA Chisel was also constrained to avoid 100% matches to any 15-base-pair window in any previously codon-optimized light J gene. The random seed in NumPy^73^ was set to 123, so the sequences will be reproduced upon reruns. A *SapI* site (“GCTCTTCC”) and one gate for ligating to the light V genes (“CCT”, encoding for proline) were added to the 5’ end of each light J sequence. 11 base pairs of random DNA (“AATACGATCAT”), a second *SapI* site (“CGAAGAGC”) and a second gate for ligating to the light Vs (“GTT”) were added to the 5’ end of the light J sequence. The kappa constant sequence, the DNA sequence in between the kappa constant and heavy V sequences, a *BsaI* site (“CGAGACC”), a six-base-pair padding sequence (“GACATT”), and an *EcoRV* site (“GATATC”) were added to the 3’ end of the light J sequence. The final light J sequences were thus constructed as follows:

GTT - *SapI* - junk DNA - *SapI* - CCT - light J coding sequence - kappa constant - sequence between kappa constant and heavy V - *BsaI* - pad - *EcoRV*

The light J sequence was ordered as a gene cloned into the pTwist Kan High Copy v2 plasmid from Twist Bioscience.

#### Analysis of Sanger sequencing of single colonies

DNA sequences were extracted from Ab1 files by aligning to a reference sequence with the same non-H3 CDR lengths on Benchling.com. Alignments were visually inspected, and base pairs labeled as “N” by the software were corrected based on the sensorgram when possible. DNA sequences were segmented into component parts (light V, light J, light constant, heavy V, CDRH3, heavy J), and each part was searched against a database of designed parts at the amino acid and DNA level. Translations were done using Biopython^75^. Sequences that could not be exactly matched were visually inspected.

#### Nanopore sequencing analysis

Although Nanopore sequencing has lower accuracy and throughput than Illumina sequencing, we felt it was appropriate to analyze the diversity of component pieces in the proof-of-concept repertoire. Additionally, Nanopore accuracy has increased to Q30 (>99.9% accuracy per base-pair) due to improved chemistry, basecalling models, and duplex sequencing^76^.

The pod5 files output from nanopore were base-called using the Dorado duplex base caller using the super accurate basecalling model. The resulting base-called reads were filtered to remove simplex reads already accounted for as duplex reads using SAMtools view^77^. The filtered reads were then demultiplexed using Dorado demux, and the adaptors were trimmed using Dorado trim.

Each read was aligned to a FASTA database of combined light V and light J sequences, a FASTA database of heavy V sequences, and a FASTA database of the one heavy J + heavy constant sequence using MiniMap2^78^ using the lr:hq preset. The alignments for each database were filtered to exclude unmapped reads and non-primary alignments using SAMtools view ^77^ was used to iterate through the alignments for each database and filter the reads based on several properties. Queries shorter than 1,300 base pairs and longer than 1,600 base pairs or with average PhRed^79^ score < 20 were filtered as were aligned sequences with lengths more than 5 bps different than their target sequences. Finally, alignments with MAPQ score < 20 were filtered. The alignments did not require a perfect match. This procedure resulted in 610,770 mapped reads from the original 1,253,324 reads.

The above pipeline does not give any information on CDR H3. To investigate this, each read was segmented using the previous alignments to extract H3 from each read. H3s with a high probability (>= 0.9) of having no errors based on the per-base pair PhRed scores were searched against a database of the designed H3s, and reads with a perfect match to a designed H3 were considered. This procedure resulted in 407,903 mapped reads, about 2/3 of the reads mapped to V genes.

#### PDB search for HEWL binders

BLASTp^80^ was used to find similar sequences to HEWL on the PDB using the pdbaa database. Hits that aligned to at least 50% of HEWL with at least 90% percent identity that are present in SAbDab^37^ were considered. Each pdb file was then loaded into PyMOL, and the antigen chain was aligned to HEWL in the mAb102 cryoEM structure. The resulting PyMOL session was visually inspected.

#### PDB search for the new Lysine-binding motif

BLASTp^80^ was used to find similar sequences to the light V gene sequence of mAb102 (IGKV1-39) on the PDB using the pdbaa database. Hits which aligned to at least 90% of IGKV1-39 with at least 90% percent identity were then cross-referenced with SAbDab^37^ to annotate the heavy and antigen chains. Each pdb file was then loaded into PyMOL, and the light chain was aligned to the light chain of mAb102. The residues in each structure closest to the two Tyrs of interest in mAb102 were determined, and only structures with Tyrs in both positions were considered. Any structures with an antigen Lys side chain within 5 Å of both Tyrs were visually inspected.

#### Data visualization

Visualization of protein structures was done using PyMOL v3.1.0. Protein surface maps were generated using the APBS^81^ plug-in in PyMOL. Data visualization was done using matplotlib v3.1.0 and seaborn v0.9.0. Data was organized using pandas v0.24.2.

### Experimental methods

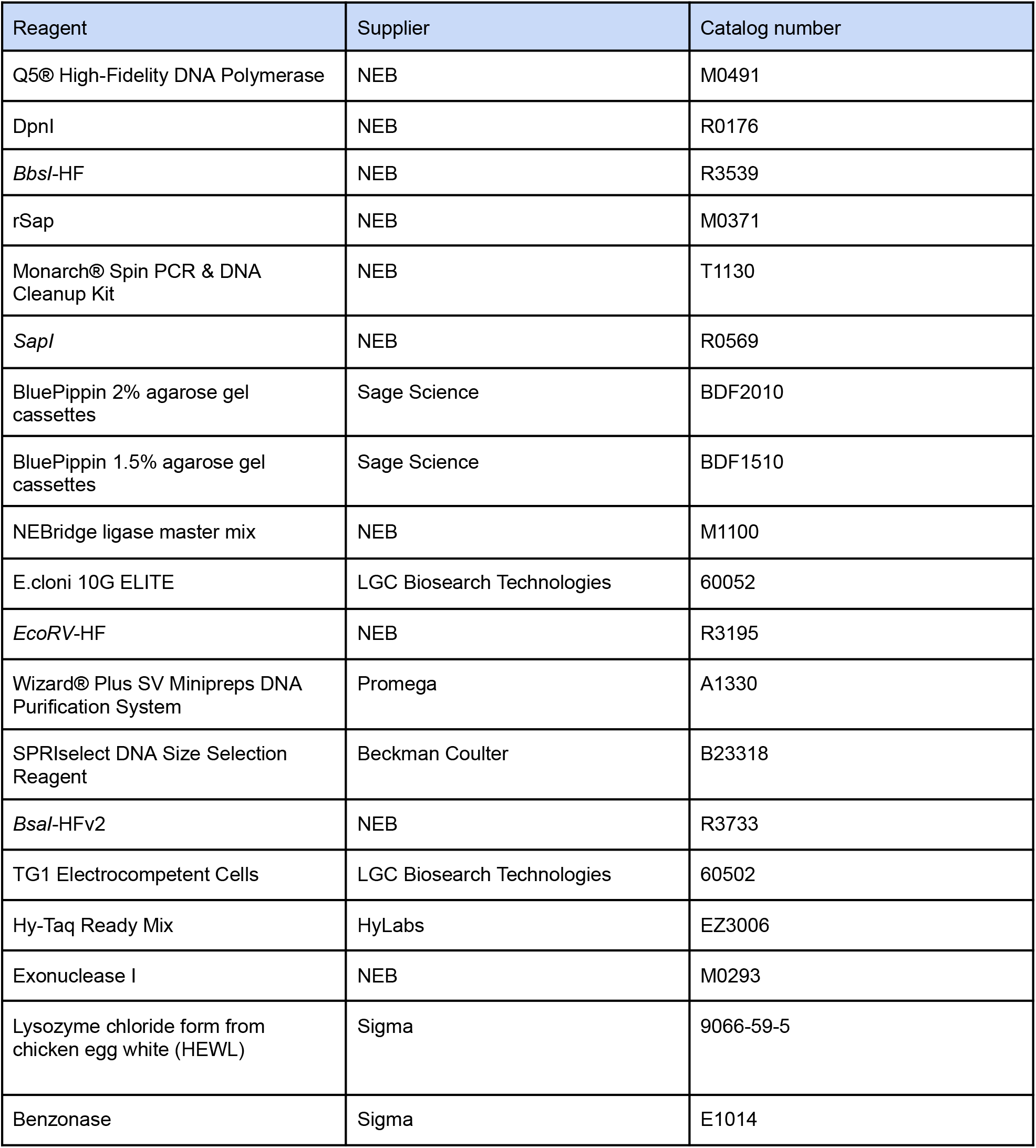
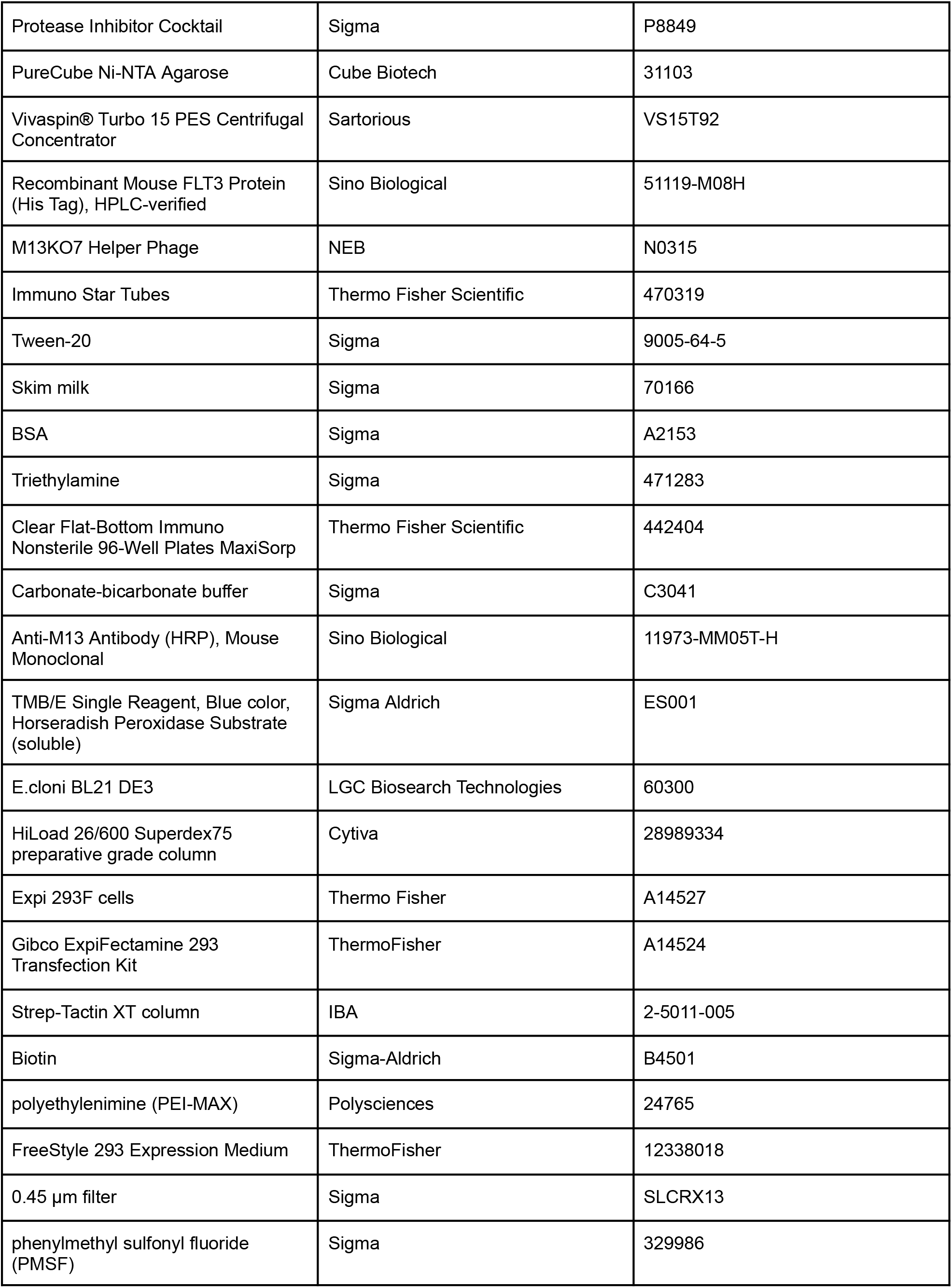
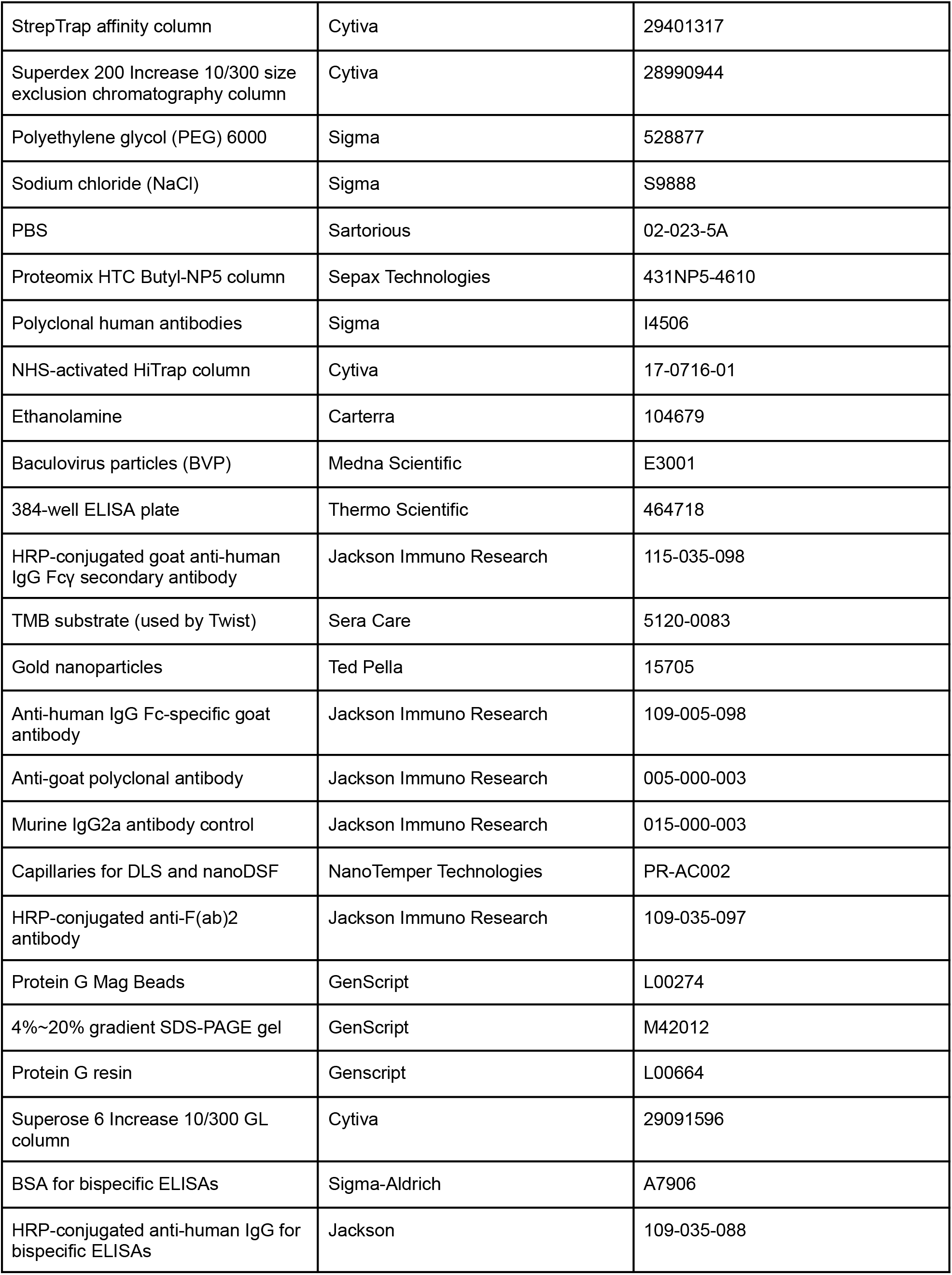
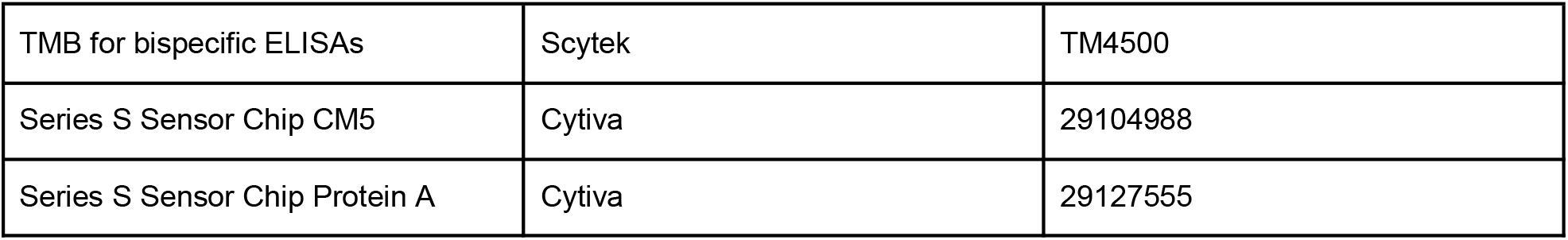

#### Preparation of a modified version of the pComb3x^44^ plasmid

The pComb3X plasmid was generously provided by the Benhar Lab (Tel Aviv University). pComb3X was modified to remove all *BsaI, SapI,* and *BbsI* sites. Additionally, an irrelevant insert was placed in the region between the OmpA leader (before the light chain) and the heavy constant region. The irrelevant insert was flanked by *BbsI* sites, so the library could be inserted using a Golden Gate^39^ reaction. The modified plasmid was PCR amplified using Q5® High-Fidelity DNA Polymerase (NEB M0491) and treated with DPNI (NEB R0176). The plasmid was then digested with *BbsI*-HF (NEB R3539). After the digestion reaction was completed, rSap (NEB M0371) was added to dephosphorylate the plasmid. The resulting product was purified using the Monarch® Spin PCR & DNA Cleanup Kit (NEB T1130), and DNA concentrations were measured using NanoDrop (Thermo Scientific). The size was verified using Tapestation (Agilent). The modified plasmid was deposited with Addgene (ID 249256).

#### Assembly of the DNA encoding for the proof-of-concept library

A detailed protocol for DNA assembly is available^40^.

Light V genes were ordered cloned into the pTwist Chlor High Copy plasmid by Twist Bioscience. The light V gene plasmids were mixed at an equimolar amount.

Light J genes were ordered as genes cloned into the pTwist Kan High Copy v2 plasmid by Twist Bioscience. The plasmids were separately digested using *SapI* (NEB R0569) to linearize the plasmids. rSap (NEB M0371) was added to each reaction to dephosphorylate the digested plasmids, reducing religation of the empty plasmids. The digested plasmids were purified using the Monarch® Spin PCR & DNA Cleanup Kit (NEB T113), and DNA concentrations were measured using NanoDrop (Thermo Scientific). The sizes were verified using Tapestation (Agilent), and the NanoDrop concentrations were corrected by the fraction of the correct size product measured by Tapestation.

The pool of light V gene plasmids was inserted into each digested light J gene plasmid in a Golden Gate^39^ reaction with *SapI* (NEB R0569) using the NEBridge ligase master mix (NEB M1100). The resulting products were purified using the Monarch® Spin PCR & DNA Cleanup Kit (NEB T1130). 5 μL of each product was transformed into E. cloni bacteria (LGC Biosearch Technologies 60052). Each transformation was titrated after one hour of recovery to verify that the entire light-chain library was transformed. The remaining bacteria were plated on 2YT plates supplemented with 50 μg/mL kanamycin and 1% glucose (2YTKG) and grown at 37°C overnight. The next day, the bacteria were taken off the plates in 5 mL 2YTKG. Five 150 μL of bacteria for each transformation were miniprepped using the Wizard® Plus SV Minipreps DNA Purification System (Promega A1330), and the minipreps for each transformation were combined. 50% glycerol was added to the remaining bacteria to a final concentration of 25% and the stocks were snap-frozen in liquid nitrogen and then stored at -80°C. The DNA concentration of the combined minipreps for each light J assembly was measured using NanoDrop (Thermo Scientific). Each light J assembly was then digested using *EcoRV*-HF (NEB R3195), and each reaction was purified using the Monarch® Spin PCR & DNA Cleanup Kit (NEB T113). DNA concentrations were measured using NanoDrop (Thermo Scientific). The products were then size-purified using a BluePippin 2% agarose gel cassette (Sage Science BDF2010). The resulting products were purified using the Monarch® Spin PCR & DNA Cleanup Kit (NEB T113), and DNA concentrations were measured using NanoDrop (Thermo Scientific). The sizes were verified using Tapestation (Agilent), and the NanoDrop concentrations were corrected by the fraction of the correct size product measured by Tapestation.

Heavy V genes were ordered cloned into the pTwist Kan High Copy v2 plasmid by Twist Bioscience. The heavy V gene plasmids were mixed at an equimolar amount. The genes were digested out of the pool of plasmids using *EcoRV*-HF (NEB R3195). The resulting product was purified using the Monarch® Spin PCR & DNA Cleanup Kit (NEB T1130), and DNA concentration was measured using NanoDrop (Thermo Scientific). The product was then size-purified using a BluePippin 2% agarose gel cassette (Sage Science, BDF2010). The resulting product was purified using the Monarch® Spin PCR & DNA Cleanup Kit (NEB T113), and DNA concentration was measured using NanoDrop (Thermo Scientific). The size was verified using Tapestation (Agilent), and the NanoDrop concentration was corrected by the fraction of the correct size product measured by Tapestation.

The heavy J gene was ordered as a gene cloned into the pTwist Kan High Copy v2 plasmid by Twist Bioscience. The gene was digested out of the plasmid using *EcoRV*-HF (NEB R3195). The resulting product was purified using the Monarch® Spin PCR & DNA Cleanup Kit (NEB T1130), and DNA concentration was measured using NanoDrop (Thermo Scientific). The product was then size-purified using a BluePippin 2% agarose gel cassette (Sage Science BDF2010). The resulting product was purified using the Monarch® Spin PCR & DNA Cleanup Kit (NEB T1130), and DNA concentration was measured using NanoDrop (Thermo Scientific). The size was verified using Tapestation (Agilent), and the NanoDrop concentration was corrected by the fraction of the correct size product measured by Tapestation.

CDR H3s were ordered as an oligo pool cloned into the pTwist Kan High Copy v2 plasmid by Twist Bioscience. The CDR H3 oligos, heavy V genes, and heavy J gene were ligated together in a Golden Gate^39^ reaction with *SapI* (NEB R0569) using the NEBridge ligase master mix (NEB M1100). After the reaction finished, additional *SapI* was added to ensure complete digestion. The resulting product was purified using the Monarch® Spin PCR & DNA Cleanup Kit (NEB T1130), and DNA concentration was measured using NanoDrop (Thermo Scientific). The size was verified using Tapestation (Agilent), and the NanoDrop concentration was corrected by the fraction of the correct size product measured by Tapestation. The heavy chain assembly was amplified by PCR using Q5® High-Fidelity DNA Polymerase (NEB M0491). The resulting product was purified with SPRIselect DNA Size Selection Reagent (Beckman Coulter B23318) using a 1.8X ratio, and DNA concentration was measured using NanoDrop (Thermo Scientific). The size was verified using Tapestation (Agilent), and the NanoDrop concentration was corrected by the fraction of the correct size product measured by Tapestation.

Heavy and light chain assemblies were ligated together in a Golden Gate^39^ reaction with *Bsa*-HFv2 (NEB R3733) using the NEBridge ligase master mix (NEB M1100), with a separate reaction for each light J. The resulting product was purified using the Monarch® Spin PCR & DNA Cleanup Kit (NEB T1130), and DNA concentration was measured using NanoDrop (Thermo Scientific). The full assemblies were amplified by PCR using Q5® High-Fidelity DNA Polymerase (NEB M0491). The resulting products were purified with SPRIselect DNA Size Selection Reagent (Beckman Coulter B23318) using a 1.8X ratio, and DNA concentrations were measured using NanoDrop (Thermo Scientific). The sizes were verified using Tapestation (Agilent), and the NanoDrop concentrations were corrected by the fraction of the correct size product measured by Tapestation. The products were then size-purified using a BluePippin 1.5% agarose gel cassette (Sage Science BDF1510). The resulting products were purified using the Monarch® Spin PCR & DNA Cleanup Kit (NEB T113), and DNA concentrations were measured using NanoDrop (Thermo Scientific). The sizes were verified using Tapestation (Agilent), and the NanoDrop concentrations were corrected by the fraction of the correct size product measured by Tapestation. The full assemblies for each light J were combined at an equimolar ratio.

#### Ligation of the library into the phagemid and transformation into bacteria

The resulting library was ligated into the pre-digested modified pComb3X^44^ vector in a Golden Gate^39^ reaction with *BbsI*-HF (NEB R3539) using the NEBridge ligase master mix (NEB M1100). The resulting product was purified using the Monarch® Spin PCR & DNA Cleanup Kit (NEB T1130). 2 μL of the product was transformed into 25 μL of commercially electrocompetent TG1 cells (LGC Biosearch Technologies 60502). The transformed bacteria were allowed to recover for 1 hour at 37°C and then pooled. 10 μL of the transformed bacteria was titrated three times, and each titration was plated three times (a total of nine replicates). The remaining bacteria were plated on five 140 mm 2YT plates supplemented with 100 μg/mL ampicillin and 1% glucose (2YTAG) and incubated at 37°C overnight. The next morning, colonies were counted on the titration plates, and the total amount of colony-forming units (cfu) in the transformed population was counted. There were >2*10^9^ cfu. 24 colonies were assayed using colony PCR to check the quality of the transformed library (see next section). The bacteria were taken off the 140 mm plates in 2YTAG. Ten 100 μL aliquots of the bacteria were miniprepped using the Wizard® Plus SV Minipreps DNA Purification System (Promega A1330). 50% glycerol was added to the remaining bacteria to a final concentration of 25% glycerol. The library was snap-frozen in liquid nitrogen and stored at -80°C.

#### Colony PCR and sequencing

Colonies were picked into 10 μL of water. Plasmid DNA was then amplified in a PCR reaction using Hy-Taq Ready Mix (HyLabs EZ3006). Product size and purity were verified using gel electrophoresis with 1% agarose gels. The remaining products were treated with 0.5 μL of Exonuclease I (NEB M0293) and rSap (NEB M0371) and then sequenced by the Life Sciences Core Facility at the Weizmann Institute of Science.

#### Production of proteinGF

A plasmid containing the DNA sequence for proteinGF^82^, an engineered version of proteinG with nM affinity to human IgG1 Fab, was generously provided by the Kossiakoff lab (U. Chicago). The plasmid contains proteinGF with an N-terminal Avi-tag followed by a TEV protease cleavage site. The Avi tag, TEV site, and proteinGF were then cloned into a pET28 plasmid with an N-terminal His-tag, followed by a bdSUMO tag. The plasmid was transformed into *E. coli* BL21 (DE3) cells (LGC Biosearch Technologies 60300). For expression, 250 mL of 2YT medium supplemented with 50 μg/ mL kanamycin was inoculated with 2.5 mL of an overnight culture produced from a single colony and grown at 37 °C until it reached an optical density (OD) 600 of 0.8. Overexpression was induced by adding 1 mM IPTG, and the cultures were grown for 20 hours at 16 °C. The cultures were collected by centrifugation for 10 min at 5,000 xg, and the pellet was frozen at −20 °C. The cells were resuspended in basic buffer (50 mM Tris-HCl pH 8, 150 mM NaCl, 10% glycerol) supplemented with 10 µg/mL lysozyme (Sigma 9066-59-5), protease-inhibitor cocktail (Sigma P8849), and benzonase (Sigma E1014). The suspension was lysed by sonication and centrifuged for 1 hour at 20,000 xg at 4 °C. The soluble fraction was filtered through a 0.45 μm vacuum filter and loaded onto a column packed with PureCube Ni-NTA Agarose (Cube Biotech 31103). The column was washed with two column volumes of MilliQ water, and then two column volumes of basic buffer before the soluble fraction was added. After the soluble fraction was added, the column was closed and incubated with shaking for 30 min at 4°C. The column was then washed with two column volumes of basic buffer supplemented with 30 mM imidazole. The protein was subjected to overnight on-column Sumo protease cleavage at 4 °C (5 μg/mL in 1.25 mL basic buffer). The next morning, the elution was collected, and another 1.25 mL basic buffer was added to the column and eluted to collect the remaining protein. The elution was concentrated using a Vivaspin Turbo centrifugal concentrator (Sartorius VS15T92) with a 3,000 kDa cutoff. The protein was purified by gel filtration (HiLoad 26/600 Superdex75 preparative grade column, Cytiva 28989334). Protein purity was assessed by SDS–PAGE. Protein concentration was determined using NanoDrop (Thermo Scientific) based on an extinction coefficient calculated using ProtParam^83^. Final aliquots were flash-frozen in liquid nitrogen and stored at –80 °C.

#### Production of SARS-CoV-2 RBD for panning

The receptor-binding domain (RBD) of SARS-CoV-2 Spike (residues 319–541), kindly provided by Florian Krammer (Mount Sinai), was modified to include a C-terminal twin Strep-tag for purification. The expression plasmid was transiently transfected into Expi 293F cells (ThermoFisher A14527) using Gibco ExpiFectamine 293 Transfection Kit (ThermoFisher A14524). After 72 hours, the culture medium (250 mL), containing the secreted RBD, was harvested. Following cell removal by centrifugation, the supernatant was dialyzed overnight at 4 °C against 50 mM Tris pH 8, 150 mM NaCl, and 1 mM EDTA (two buffer exchanges). The dialyzed medium was filtered and loaded onto a 5 mL Strep-Tactin XT column (IBA 2-5011-005) pre-equilibrated with the same buffer. After washing with equilibration buffer, the RBD was eluted using 50 mM biotin (Sigma-Aldrich B4501) in the same buffer. Elution fractions containing RBD were pooled and dialyzed overnight against PBS. Protein identity and purity were confirmed by SDS-PAGE followed by Coomassie staining. Final aliquots were flash-frozen in liquid nitrogen and stored at –80 °C.

#### Production of SARS-CoV-2 RBD for ELISAs

The receptor-binding domain (RBD) of SARS-CoV-2 Spike (residues 319–541) was produced by GenScript with a C-terminal twin Strep-tag for purification. The amino acid sequence was back-translated into DNA and codon optimized for expression in HEK cells using GeneSmart. The DNA was synthesized and cloned into the pcDNA3.4 vector. The plasmid was maxiprepped to yield transfection-grade plasmids using GenScript Proprietary in-house columns and reagents. The plasmid was then transiently transfected into a suspension HD 293F cell culture. Briefly, suspension HD 293F cells were grown in serum-free expression medium containing GenScript proprietary in-house buffers and supplements in Erlenmeyer Flasks at 37°C with 8% CO_2_ on an orbital shaker. One day before transfection, the cells were seeded at an appropriate density in an Erlenmeyer flask. On the day of transfection, DNA and transfection reagent were mixed at an optimal ratio and then added to the flask with cells ready for transfection. The cell culture supernatants were collected by centrifugation at 3,000 rpm for 30 min on day 6. The supernatants were filtered and loaded onto an affinity purification column. After washing and eluting with appropriate buffers, the eluted fractions were collected, and buffer exchanged to 1X PBS pH 7.2. The purified protein was analyzed by SDS-PAGE to determine the molecular weight and purity. Each sample was run in reducing sample buffer (60 mM Tris-HCl, 2% SDS, 6% Glycerol, 0.1% bromophenol blue, 50 mM DTT, pH 6.8) and non-reducing sample buffer (50 mM Tris-HCl, 1% SDS, 6% Glycerol, 0.1% bromophenol blue, 50 mM NEM, pH 6.8). The endotoxin level of protein was also detected using a Kinetic Turbidimetric Assay (Bioendo). The concentration was determined by measuring A280 using a NanoDrop Ultra (ThermoScientific).

#### Production of Lujo spike

DNA encoding for the Lujo spike ectodomain was ordered from Synbio Technologies into the pcDNA3.1 plasmid. A trimerization domain (GCN4) and a Strep tag (WSHPQFEK) were appended to the C-terminus of the Lujo spike.

The expression plasmid was transfected into HEK293F cells at ∼1.0 × 10⁶ cells/mL using 40 kDa polyethylenimine (PEI-MAX) (Polysciences 24765) at a ratio of 1:3 (DNA: PEI solution). The cells were grown in FreeStyle 293 Expression Medium (ThermoFisher 12338018) and incubated at 37°C with 8% CO_2_. The media was collected 6 days post-transfection by centrifugation at 500 × g for 10 min at 4°C. The supernatant was transferred to another tube and centrifuged again at 10,000 × g for 20 min at 4°C. The supernatant was filtered through a 0.45 µm filter (Sigma SLCRX13) with 0.02% sodium azide and 100 μM phenylmethyl sulfonyl fluoride (PMSF; Sigma 329986). The soluble LUJV ectodomain was purified using a StrepTrap (Cytiva 29401317) affinity column and Superdex 200 Increase 10/300 size exclusion chromatography (Cytiva 28990944). Final aliquots were flash-frozen in liquid nitrogen and stored at –80 °C.

#### Production of phage from library glycerol stocks

A 1 mL glycerol stock (OD600 of 32) of the library was added to 200 mL prewarmed 2YT supplemented with 100 μg/mL ampicillin and 1% glucose (2YTAG). The bacteria were grown at 37°C with 250 rpm shaking to an OD600 of 0.4-0.6. 1 mL (1 * 10^11^ pfu) of M13KO7 helper phage (NEB N0315) was added to 50 mL of bacteria. The bacteria were incubated at 37°C for 30 min without shaking and then 30 min with 250 rpm shaking. The bacteria were centrifuged for 10 min at 3,300 xg, resuspended in 100 mL 2YTAG supplemented with 50 μg/mL kanamycin (2YTAGK), and incubated overnight at 30°C with 250 rpm shaking. The following day, the bacteria were centrifuged for 30 min at 3,300 xg at 4°C, and the supernatant was filtered through a 0.22 μm vacuum filter. 20 mL PEG/NaCl (20% PEG 6000 [Sigma 528877], 2.5M NaCl [Sigma S9888]) was added to the supernatant, and the mixture was incubated on ice for one hour. The mixture was centrifuged for 30 min at 3,300 xg at 4°C. The supernatant was discarded, and the remainder was centrifuged for 3 min at 3,300 xg at 4°C. The pellet was resuspended in 4 mL 1X PBS (Sartorius 02-023-5A) and centrifuged for 10 min at 11,600 xg. The supernatant was then moved to a fresh tube. 10 μL of the phage was diluted in 1X PBS in ten serial dilutions of 10X, and 10 μL of each dilution was used to infect 90 μL of TG1 OD600 0.4-0.6. The infected bacteria were incubated at 37°C for 1 hour without shaking. 10 μL of each infection was plated on 2YTAG plates in replicates. The following day, the titer of the phage was calculated. The phage in PBS was either stored at 4°C if used the next day, or 50% glycerol was added to a final concentration of 25% glycerol, and the phage was snap frozen in liquid nitrogen and stored at -80°C.

#### Phage display of designed library against RBD, HEWL, Lujo, and FLT3

RBD and Lujo spike were produced as described above. HEWL (Sigma 9066-59-5) and his-tagged mouse FLT3 (Sino Biological 11973-MM05T-H) were purchased.

In all panning campaigns, the antigen was immobilized on immunotubes (ThermoFisher 470319) in 1X PBS at 10 μg/mL in a volume of 4 mL overnight at 4°C. Antigens thawed from the freezer were centrifuged for 5 min at 10,000 xg at 4°C before coating. Immunotubes were washed 3 times with 5 mL 1X PBS supplemented with 0.05% Tween-20 (PBS-T, Sigma 9005-64-5). Immunotubes were then blocked for two hours with blocking buffer (either 2% skim milk [Sigma 70166] or 3% BSA [Sigma A2153] in PBS-T) at room temperature. In the meantime, phage were blocked rotating at room temperature at a concentration of 0.25 * 10^13^ pfu/mL in blocking buffer for two hours. The immunotubes were washed 3 times with 5 mL PBS-T, and 1*10^13^ pfu of blocked phage per antigen was added to the blocked immunotubes in a total volume of 4 mL. The phage were incubated for 1 hour rotating at room temperature, and then one hour standing at room temperature. The phage were washed 10 times with 5 mL PBS-T and then 10 times with 5 mL PBS. 1 mL 100 mM triethylamine (Sigma 471283) was added to elute the phage, and the phage were incubated for 30 min with the triethylamine at room temperature, rotating. The 1 mL elution was immediately added to 500 μL 1M Tris pH 7.4, mixed, and left on ice. 1 mL of the eluted phage was used to infect 9 mL of TG1 at OD600 0.4-0.6. The infected bacteria were incubated at 37°C without shaking for 30 min and then at 37°C with shaking at 250 rpm. The bacteria were centrifuged at 3,300 xg for 10 min, resuspended in 500 μL 2YT, and plated on 140 mm 2YTAG plates. Additionally, the eluted phage was serially diluted in 10X dilutions, and 10 μL of each dilution was used to infect 90 μL of TG1 at OD600 0.4-0.6. The infected bacteria were incubated at 37°C without shaking for 1 hour. 10 μL of each infection was plated on 2YTAG plates in replicates.

The following day, the titer of the phage was calculated. Additionally, the bacteria were taken off the 140 mm 2YTAG plates with 3-5 mL 2YTAG. Several 150 μL aliquots were miniprepped using the Wizard® Plus SV Minipreps DNA Purification System (Promega A1330). 100 μL of bacteria were added to 100 mL 2YTAG and grown at 37°C with shaking at 250 rpm to an OD600 of 0.4-0.6. 50% glycerol was added to the remaining bacteria to a final concentration of 25%, and the bacteria were snap frozen in liquid nitrogen and stored at -80°C. 200 μL of M13KO7 helper (NEB N0315) phage was added to 10 mL of the bacteria, and the same protocol (with volumes adjusted) for producing the library phage was then followed to produce input phage for the next selection round.

For the first panning campaign against HEWL and RBD, there were three panning rounds, and the blocking buffer was alternated between each round (skim milk, then BSA, then skim milk). For the panning campaigns against Lujo, FLT3, and the repeat of RBD, there were two panning rounds, and skim milk was used as the blocking buffer for each. After each round, polyclonal ELISAs were performed to monitor the binding to the antigen and an off-target antigen.

#### Polyclonal phage ELISAs

Antigens were coated on 96-well MaxiSorp plates (ThermoFisher 442404) at 2 μg/mL in carbonate-bicarbonate buffer (CBC, Sigma C3041) overnight at 4°C in a volume of 50 μL. Wells were washed 3 times with 200 uL PBS-T and blocked with 2% skim milk (Sigma 70166) in PBS-T for 1 hour at room temperature. Phage were blocked at a 1:10 dilution (typically ∼10^12^ cfu/mL) with 2% skim milk in PBS-T, rotating at room temperature for one hour. Wells were washed 3 times with 200 μL PBS-T, and 50 μL blocked phage were added at several dilutions. After one hour, wells were washed 3 times with 200 μL PBS-T, and 50 μL HRP-conjugated anti-M13 antibody (Sino Biological 11973-MM05T-H) diluted to 250 ng/mL in 2% skim milk in PBS-T was added to each well. After one hour, wells were washed 3 times with 200 μL PBS-T, and 50 μL tetramethylbenzidine (TMB, Sigma ES001) was added. Once the signal saturated, 25 μL of 1M H_2_SO_4_ was added to quench the reaction, and OD450 was measured. For each sample, background binding to a well coated only with CBC was subtracted.

#### Monoclonal phage ELISAs

Several dilutions of glycerol stocks from the output of the final round of panning were plated on 2YTAG plates. Single colonies were picked into 100 μL 2YTAG in 96-well plates. Bacteria were incubated in a 96-well plate shaking incubator at 37°C overnight. The next morning, 2 μL from each well was diluted into 198 μL fresh 2YTAG in a round-bottom plate. The bacteria were incubated in a 96-well plate shaking incubator at 37°C for 2-3 hours. 5 μL of each well was plated on 2YTAG plates and grown overnight at 37°C. 50% glycerol was added to a final concentration of 25% glycerol to the remaining bacteria in each well, and the bacteria were stored at -80°C. After 2-3 hours, 10^9^ helper phage (NEB N0315) were added to each well in 25 uL 2YTAG. The infected bacteria were incubated in a 96-well plate shaking incubator at 37°C for 1 hour. The plate was centrifuged at 1,800 xg for 10 min, and the pellets were resuspended in 2YTAK. The bacteria were incubated overnight in a 96-well plate shaking incubator at 30°C. The following morning, the supernatant was mixed 1:1 with 2% skim milk (Sigma 70166) in PBS-T and used as input to an ELISA (same protocol as for polyclonal ELISAs). TMB (Sigma ES001) was allowed to develop for 10 minutes. Signals were then normalized by their respective binding to wells coated with proteinGF. The colonies were assayed by colony PCR and sequenced (see above colony PCR protocol).

Antibody production and developability testing overview:

Nine designed antibodies were selected for developability testing. Sequences were codon optimized for mammalian cell expression, and antibodies were expressed using an Expi293 expression system (ThermoFisher) by Twist Bioscience in 8mL cell cultures and purified with Protein A.

#### Developability testing overview

Each test antibody was characterized by Twist Bioscience in the following assays: analytical size-exclusion chromatography (aSEC), hydrophobic interaction chromatography (HIC), cross-interaction chromatography (CIC), baculovirus particle (BVP) ELISA, dynamic light scattering (DLS), and nano differential scanning flourimetry (nanoDSF). The antibodies were characterized alongside twelve control antibodies previously evaluated for developability characteristics (adapted from a previous paper charting the developability of therapeutic antibodies^2^).

#### Analytical size-exclusion chromatography (aSEC)

Analytical SEC was performed by Twist Bioscience on an Agilent 1260 Infinity II HPLC system equipped with an AdvanceBio SEC HPLC column 2.7mm (Agilent). Purified antibodies (10 µg) were eluted with 1x PBS, pH 7.4, at a flow rate of 0.35 mL/min, while monitoring at 280nm. Chromatogram traces were collected and analyzed using the Shimadzu LabSolutions software. Based on a single experiment.

#### Hydrophobic interaction chromatography (HIC)

HIC testing of the purified antibodies was performed by Twist Bioscience. 5 µg of antibody was run over a Proteomix HTC Butyl-NP5 column, 4.6 × 100 mm (Sepax Technologies 431NP5-4610) at 1 mL/min for a 20 min gradient. The Agilent 1260 Infinity II HPLC system monitored absorbance at 280 nm. Buffers used were 1.8 M ammonium sulfate and 100mM sodium phosphate at pH 6.5 (Mobile Phase A) and 100mM sodium phosphate at pH 6.5 (Mobile Phase B). Chromatogram traces were collected and analyzed with Agilent Open Lab CDS software. Based on a single experiment.

#### Cross-interaction chromatography (CIC)

CIC testing of the purified antibodies was performed by Twist Bioscience. A CIC column was prepared by covalently coupling ∼30 mg of polyclonal human antibodies (Sigma I4506) to a 1-mL NHS-activated HiTrap column (Cytiva 17-0716-01) and subsequently quenched with ethanolamine (Carterra 104679). For each analysis, ∼2.5 μg of antibody was loaded, with 1X PBS, pH 7.4 as the mobile phase at 0.1 mL/min on an Agilent 1260 Infinity II HPLC system. Chromatogram traces were collected and analyzed using Agilent Open Lab CDS software. Based on a single experiment.

#### Baculovirus ELISA (BVP ELISA)

BVP ELISA testing of the antibodies was performed by Twist Bioscience. In brief, baculovirus particles (Medna Scientific E3001) working solution were prepared with 50 mM sodium carbonate buffer (pH 9.6) in a 1:200 dilution and coated onto a 384-well ELISA plate (Thermo Scientific 464718). The following day, unbound particles were removed, and assays were performed at room temperature. Wells were blocked with PBS containing 2% BSA for 1 hr, washed with 1X PBS, pH 7.4 3 times, and incubated with test antibodies (100 μg/mL, 2.5 ug) in quadruplicates for 1 h. After washing 5 times with 1X PBS, pH 7.4, HRP-conjugated goat anti-human IgG Fcγ secondary antibody (Jackson Immuno Research 115-035-098) was added for 1 h, washed 5 times with 1X PBS, pH 7.4 followed by TMB substrate (Sera Care 5120-0083) incubation for 10–15 min. The reaction was stopped with 2 M sulfuric acid, and absorbance was read at 450 nm using an Agilent Biotech Epoch Microplate Spectrophotometer reader. Fold-change was calculated by normalizing the OD of the measured well to that of a well containing secondary antibody but not the tested antibody. Measurements were performed in quadruplicate.

#### Affinity Capture Self-Interaction Nanoparticle Spectroscopy (AC-SINS)

AC-SINS testing of the antibodies was performed by Twist Bioscience. Gold nanoparticles (Ted Pella 15705) were mixed with 80% anti-human IgG Fc-specific goat antibody (Jackson Immuno Research 109-005-098) and 20% anti-goat polyclonal antibody (Jackson Immuno Research 005-000-003) at room temperature. Both antibodies were buffer exchanged to 43 mM Sodium Acetate, pH 4.3, prior to mixing. Test samples were incubated with the coated particles for 2 h at room temperature in quadruplicate. Absorbance spectra were acquired using an Agilent Biotech Epoch Microplate Spectrophotometer, and maximum absorption wavelength (λmax) values were determined. A shift in peak absorbance wavelength was determined by subtracting the λmax from an isotype negative control (muIgG2a) (Jackson Immuno Research 015-000-003). Final normalized values are reported as Δλmax. Measurements were performed in quadruplicate.

#### Dynamic Light Scattering (DLS)

DLS testing of the antibodies was performed by Twist Bioscience. Protein samples (∼10 µL, 0.5 mg/mL) were loaded into standard capillaries (NanoTemper Technologies PR-AC002) and analyzed on a Prometheus Panta instrument. Sample information, including protein identity, buffer composition, and concentration, was recorded in the PR Panta Analysis software (v1.9) prior to measurement. 10 acquisitions were performed at 20°C, and distribution profiles were analyzed in PR.Panta Analysis v1.9 to determine polydispersity index (PDI) and hydrodynamic radius (RI). Each measurement was performed in duplicate.

#### Nano Differential Scanning Fluorimetry (nanoDSF)

NanoDSF testing of the antibodies was performed by Twist Bioscience. Protein samples (∼10 µL, 0.5 mg/mL) were loaded into standard capillaries (NanoTemper Technologies PR-AC002). Excitation parameters were optimized by discovery scans, after which thermal unfolding was assessed from 15 °C to 95 °C at a 1 °C/min ramp rate. Traces and parameters were determined using PR.Panta Analysis v1.9. Fluorescence intensity ratios (F350/F330) were plotted against temperature, with Tonset defined as the initial deviation from baseline and Tm’s determined from the inflection point of the first derivative. Each measurement was performed in duplicate.

#### Antigen ELISAs with IgG

Antigens were coated on 96-well MaxiSorp plates (ThermoFisher 442404) at 2 μg/mL in carbonate-bicarbonate buffer (CBC, Sigma C3041) overnight at 4°C in a volume of 50 μL. Wells were washed 3 times with 200 uL PBS-T and blocked with 2% skim milk (Sigma 70166) in PBS-T for 1 hour at room temperature. Wells were washed 3 times with 200 μL PBS-T, and 50 μL of antibody dilutions in 2% skim milk in PBS-T were added to each well. After one hour, wells were washed 3 times with 200 μL PBS-T, and 50 μL HRP-conjugated anti-F(ab)_2_ antibody (Jackson Immuno Research 109-035-097) diluted to 400 ng/mL in 2% skim milk in PBS-T was added to each well. After one hour, wells were washed 3 times with 200 μL PBS-T, and 50 μL tetramethylbenzidine (TMB, Sigma ES001) was added. Once the signal saturated, 25 μL of 1M H_2_SO_4_ was added to quench the reaction, and OD450 was measured. Curves were fit using Nonlinear regression (curve fit) in Graphpad Prism v10.4.2. Experiments were performed in duplicate.

For the specificity heat map, ELISAs were completed as described above with 100nM, 10nM, and 1nM of each antibody in duplicate. Area under the curve (AUC) was computed using Graphpad Prism v10.4.2. A control antibody was included on each plate for comparison. Experiments were performed in duplicate.

#### mAb102 and 601 Fab production

The antibodies were produced by GenScript. The amino acid sequences for the light and heavy chains were back-translated into DNA and codon optimized for expression in CHO cells using GeneSmart. The DNA was synthesized and cloned into an optimized human IgG1 Fab expression vector for both the light and heavy chains. The plasmids were maxiprepped to yield transfection-grade plasmids using GenScript Proprietary in-house columns and reagents. The plasmids were then transiently co-transfected at a 1:1 ratio into suspension CHO-Express cell cultures. Briefly, suspension CHO cells were grown in serum-free expression medium containing GenScript proprietary in-house buffers and supplements in Erlenmeyer Flasks at 37°C with 5% CO2 on an orbital shaker. One day before transfection, the cells were seeded at 3-6x106 cells/mL in an Erlenmeyer flask. On the day of transfection, DNA and transfection reagent were mixed at an optimal ratio and then added to the flask with cells ready for transfection. The cell culture supernatants were collected by centrifugation at 3,000 rpm for 30 min when the cell viability was less than 50%. The supernatants were filtered and loaded onto Protein G Mag Beads (GenScript L00274) and incubated for the appropriate time. The beads were washed and eluted using GenScript Proprietary in-house columns and reagents. The eluted fractions were pooled and buffer exchanged to 1X PBS pH 7.2. The purified protein was analyzed by SDS-PAGE analysis using a 4%∼20% gradient SDS-PAGE gel (GenScript M42012) to determine the molecular weight and purity. Each sample was run in reducing sample buffer (60 mM Tris-HCl, 2% SDS, 6% Glycerol, 0.1% bromophenol blue, 50 mM DTT, pH 6.8) and non-reducing sample buffer (50 mM Tris-HCl, 1% SDS, 6% Glycerol, 0.1% bromophenol blue, 50 mM NEM, pH 6.8). The endotoxin level of protein was also detected using a Kinetic Turbidimetric Assay (Bioendo). The concentration was determined by measuring A280 using a NanoDrop Ultra (ThermoScientific).

#### Bispecific antibody production

Bispecific antibodies were generated in a human IgG1 format using CrossMab and knob-into-hole heterodimerization technologies and transiently expressed in Expi293 cells. The resulting antibodies were purified from culture supernatants by Protein G affinity chromatography (GenScript L00664), buffer-exchanged into PBS, sterile filtered, and stored at 4°C until use.

#### Size-exclusion chromatography (SEC) of bispecific antibodies

Purified bispecific antibodies were analyzed by size-exclusion chromatography (SEC) using a Superose 6 Increase 10/300 GL column (Cytiva 29091596) connected to an ÄKTA Pure chromatography system. Elution profiles were monitored by UV absorbance at 280 nm to assess antibody purity and aggregation. Chromatogram traces were collected and analyzed using the Shimadzu LabSolutions software. Based on a single experiment.

#### Antigen ELISAs with bispecific IgG

Binding of bispecific antibodies to FLT3 was evaluated by ELISA. High-binding 96-well plates were coated overnight at 4°C with recombinant FLT3 (1 μg/mL, Sino 51119-M08H). Plates were blocked with 2% BSA (Sigma-Aldrich A7906) in PBS and incubated with serial dilutions of bispecific antibodies. Bound antibodies were detected using HRP-conjugated anti-human IgG (Jackson 109-035-088), followed by TMB (Scytek TM4500) substrate development. Absorbance was measured at 450 nm using a microplate reader. The experiment was performed in duplicates.

#### Surface plasmon resonance (SPR)

SPR experiments were carried out on a Biacore 8K instrument (Cytiva) at 25 °C with PBST (1X PBS + 0.05% Tween-20). Each experiment was based on a single measurement.

For binding analysis of antibodies against monovalent targets, each antibody was immobilized to a flow cell of a protein A sensor chip (Cytiva 29127555). A reference flow cell for each channel was not loaded with antibody. Approximately 150-250 response units of each anti-HEWL and ant-FLT3 antibody or approximately 500 response units of each anti-RBD antibody were immobilized. Samples of different antigen (HEWL: Sigma 9066-59-5, RBD: produced by Genscript, mFLT3: Sino 51119-M08H) concentrations were injected over the surface and the chip was washed with PBST. Concentrations used for each antigen can be found in Supp. Table 13. Flow rates and times for each step can be found in Supp. Table 17. After each injection, surface regeneration was performed with two 30-second injections of 10 mM glycine pH 1.5 at a flow rate of 10 μL/min, and reached the same baseline. The sensogram for each reference flow cell was subtracted from the sensogram for the flow cell with antibody from the same channel. The acquired sensograms were analysed using the Biacore Insight Evaluation Software (v.4.0.8.19879), with five concentrations selected for fitting. Dissociations were manually trimmed to remove flat baselines. Full multi-cycle kinetic fits to a 1:1 binding model were performed with global fitting of kon, Rmax, and tc with standard initial values. RI and drift were set to a constant 0.

Because the Lujo spike is a trimer, the spike protein was immobilized on a CM5 chip (Cytiva) and mAb601 Fab was flown over the chip. The Lujo protein lost binding with regeneration, so single-cycle kinetics analysis was performed. The Lujo spike protein was immobilized to a flow cell of a CM5 chip (Cytiva 29104988) using Amine Coupling Kit (Cytiva BR100633) at a concentration of 10 μg/mL in 150mM Sodium Acetate pH 4.6 for 45 sec at a flow rate of 10 μL/min. A reference flow cell for each channel was not loaded with protein. Approximately 450 response units of the Lujo spike were immobilized. Samples with different concentrations of mAb601 Fab (produced by Genscript) were injected over the surface at a flow rate of 30 μL/min for 100 s, and the chip was washed with PBST for 900 sec at a flow rate of 30 μL/min. Concentrations of Fab601 used can be found in Supp. Table 13. The sensogram for the reference flow cell was subtracted from the sensogram for the flow cell coated with Lujo from the same channel. The acquired sensograms were analysed using the Biacore Insight Evaluation Software (v.4.0.8.19879). A single-cycle kinetic fit to a 1:1 binding model was performed with global fitting of kon, Rmax, and tc with standard initial values. RI and drift were set to a constant 0.

#### Cryo-EM data collection and analysis

3 µl of a 5 mg/mL mix of mAb102 and HEWL (Sigma 9066-59-5) at a 1:1 mass ratio was applied to glow-discharged (120 s, 15 mA; Pelco easiGlow, Ted Pella) UltrAuFoil R1.2/1.3 grids (Quantifoil Micro Tools GmbH) and plunge frozen using Vitrobot Mark IV (Thermo Fisher Scientific) at 22°C, 100% humidity, and 4 s blot time. Cryo-EM data were collected using Talos Arctica G2 microscope operated at 200 kV using Falcon 4i direct detector (Thermo Fisher Scientific) at a nominal magnification of ×190,000, corresponding to a physical pixel size 0.731 Å. 12,149 .eer movies were recorded using EPU 3.14 suite (Thermo Fisher Scientific) at a dose rate of 8.4 e/pix/s and total exposure time of 3.9 s, resulting in accumulated dose of 60 e/Å2. The nominal defocus range was −1 to −2.5 μm.

Data processing was carried out using CryoSPARC v4.7.1 (10.1038/nmeth.4169). All movies were subjected to Patch Motion correction, followed by Patch CTF estimation. Movies with thick ice or poor CTF fits were discarded. The processing scheme is outlined in Supp. Fig. 19. Initial particle picking was performed using the Blob picker tool. Extracted particles were iteratively cleared by 2D classification, ab initio, and heterogeneous refinements to produce initial models and 2D templates. Then 17,250,033 particles were selected using the Template picker tool and extracted with a 512-pixel box size, Fourier-cropped to a 256-pixel box (1.462 Å/pix). The dataset was cleaned by iterative 2D classifications, heterogeneous and non-uniform refinements, and 3D classifications. After removing duplicates and rebalancing orientations, the resulting 175,447 particles were subjected to Global and Local CTF refinement and Reference-Based motion correction. The final map was obtained using non-uniform refinement without symmetry imposed and was filtered to the local resolution before atomic model refinement.

#### Cryo-EM Model building, refinement, and analysis

The initial model for the Fab was generated using the AlphaFold3^84^ web server, and the initial model for HEWL was derived from the PDB (PDB ID: 1MLC). These initial models were fitted into the EM map using ChimeraX^85^ and were refined using manual fitting in Coot^86^ and real-space refinement in Phenix^87^. Structural analysis and representation were done using PyMOL and ChimeraX.

## Contributions

A.T. and S.J.F. conceived the idea. A.T. and S.J.F. wrote the paper, with input from all authors. A.T. developed the design algorithm and designed all DNA and amino acid sequences. A.T. developed the DNA assembly protocol and assembled the proof-of-concept library together with S.G.. A.T. performed phage display panning, and R.W. and S.G. performed phage ELISAs. T.U. and S.A. expressed and purified the RBD protein. S.B. expressed and purified the Lujo spike protein and performed the Lujo-related assays. S.B., R.D., A.T., R.W., and S.J.F. analyzed the Lujo-related data. R.W. and S.G. performed the IgG ELISAs, and R.W., S.G., A.T., and S.J.F. analyzed the data. A.T. performed the deep sequencing and analyzed the data. M.G. analyzed the SEC data and provided experimental support. E.H.Y. and R.W. performed the SPR experiments. I.C. prepared the samples for cryo-EM analysis. R.K. performed the cryo-EM experiment, and R.K., A.T., I.C., R.D., and S.J.F. analyzed the data. M.F. performed the bispecific antibody experiments, and M.F., R.D., A.T., S.J.F, and M.G. analyzed the data. S.J.F. was responsible for funding acquisition.

## Acknowledgments

We thank the Benhar Lab (Tel Aviv University) for generously providing the pComb3X plasmid, the Kossiakoff Lab (University of Chicago) for a plasmid encoding for proteinGF, and Florian Krammer (Mount Sinai) for a plasmid encoding the receptor-binding domain (RBD) of SARS-CoV-2 Spike. We thank Ahuva Nissim (Queen Mary University of London and Weizmann Institute of Science) and Adva Mechaly (Israel Institute for Biological Research) for invaluable advice on phage display and library generation and critical discussions on the work. We thank Dina Listov, Olga Khersonsky, Lucas Krauss (Weizmann Institute of Science), Peter Tessier (University of Michigan), Gilad Kaplan, Mark Austin, Daniela Zanatta, Thomas M. Moon (AstraZeneca), Moshe Ben-David (Ukko), and Hezi Sasson (Biolojic Design) for critical discussions. A.T. was supported by Teva Pharmaceutical Industries Ltd as part of the Israeli National Forum for BioInnovators (NFBI) and by the Azrieli Foundation under the Azrieli Fellows Program. The study was supported by a European Research Council Advanced Grant (101140394) to S.J.F.

## Data availability

All data generated or analyzed during this study are included in this published article and its supplementary files, with the following exceptions: Human antibody germline sequences were taken from the IMGT reference database (https://www.imgt.org/vquest/refseqh.html), sequences for therapeutic controls were taken from the TheraSAbDab database (http://opig.stats.ox.ac.uk/webapps/newsabdab/therasabdab/search), and antibody structures were taken from the Protein Data Bank (https://www.rcsb.org/). The bam files from the deep sequencing of the unselected repertoire have been deposited in the Figshare database under accession code 32968685. The raw pod5 files can be provided upon reasonable request. Heavy and light V genes used in the full designed CADAbRe repertoire and their corresponding amino acid sequences are available in Supp. Table 3. Heavy and light V and J genes in the assembled proof-of-concept CADAbRe repertoire and their corresponding amino acid and DNA sequences are available in Supp. Tables 4-7. Amino acid and DNA sequences of designed CDR H3s in the assembled proof-of-concept CADAbRe repertoire are available in Supp. Table 9. Coordinate files and experimental density maps have been deposited at the Protein Data Bank (PDB) and Electron Microscopy Data Bank (EMDB) under accession codes 9U0E and EMD-56486, respectively. The proof-of-concept repertoire will be made available to academics upon request. Source data are provided with this paper.

## Code Availability

RosettaScripts^66^ XMLs for running CADAbRe are available at https://github.com/Fleishman-Lab/CADAbRe_public.

## References

1. EARA/EFPIA Response to EURL ECVAM Recommendation on Non-Animal-Derived Antibodies.

2. Jain, T. et al. Biophysical properties of the clinical-stage antibody landscape. Proc. Natl. Acad. Sci. U. S. A. 114, 944–949 (2017).

3. Ferrara, F. et al. A next-generation Fab library platform directly yielding drug-like antibodies with high affinity, diversity, and developability. MAbs 16, 2394230 (2024).

4. Azevedo Reis Teixeira, A., et al. Drug-like antibodies with high affinity, diversity and developability directly from next-generation antibody libraries. MAbs 13, 1980942 (2021).

5. Kothiwal, D., et al. High-throughput machine learning-aided antibody discovery for cell surface antigens. bioRxiv (2025) doi:10.1101/2025.05.15.650607.

6. Putyrski, M. et al. Pioneer: a synthetic human antibody phage display library for rapid therapeutic lead generation. MAbs 17, 2543769 (2025).

7. Publications Office of the European Union. *EURL ECVAM Recommendation on Non-Animal-Derived Antibodies*. (Publications Office of the European Union, 2020).

8. Crescioli, S. et al. Antibodies to watch in 2025. MAbs 17, 2443538 (2025).

9. Gordon, G. L., Raybould, M. I. J., Wong, A. & Deane, C. M. Prospects for the computational humanization of antibodies and nanobodies. Front. Immunol. 15, 1399438 (2024).

10. Winter, G. & Milstein, C. Man-made antibodies. Nature 349, 293–299 (1991).

11. Bradbury, A. R. M., Sidhu, S., Dübel, S. & McCafferty, J. Beyond natural antibodies: the power of in vitro display technologies. Nat. Biotechnol. 29, 245–254 (2011).

12. Prassler, J. et al. HuCAL PLATINUM, a synthetic Fab library optimized for sequence diversity and superior performance in mammalian expression systems. J. Mol. Biol. 413, 261–278 (2011).

13. Valadon, P. et al. ALTHEA Gold Libraries^TM^: antibody libraries for therapeutic antibody discovery. MAbs 11, 516–531 (2019).

14. Tiller, T. et al. A fully synthetic human Fab antibody library based on fixed VH/VL framework pairings with favorable biophysical properties. MAbs 5, 445–470 (2013).

15. Gray, A. et al. Animal-free alternatives and the antibody iceberg. Nat. Biotechnol. 38, 1234–1239 (2020).

16. Jain, T. et al. Biophysical properties of the clinical-stage antibody landscape. Proc. Natl. Acad. Sci. U. S. A. 114, 944–949 (2017).

17. Makowski, E. K. et al. Optimization of therapeutic antibodies for reduced self-association and non-specific binding via interpretable machine learning. *Nat*. Biomed. Eng. 8, 45–56 (2024).

18. EARA/EFPIA Response to EURL ECVAM Recommendation on Non-Animal-Derived Antibodies.

19. Foote, J. & Winter, G. Antibody framework residues affecting the conformation of the hypervariable loops. J. Mol. Biol. 224, 487–499 (1992).

20. Makabe, K. et al. Thermodynamic consequences of mutations in vernier zone residues of a humanized anti-human epidermal growth factor receptor murine antibody, 528. J. Biol. Chem. 283, 1156–1166 (2008).

21. Tennenhouse, A. et al. Computational optimization of antibody humanness and stability by systematic energy-based ranking. *Nat*. Biomed. Eng. 8, 30–44 (2024).

22. Almagro, J. C., Pedraza-Escalona, M., Arrieta, H. I. & Pérez-Tapia, S. M. Phage display libraries for antibody therapeutic discovery and development. Antibodies (Basel) 8, 44 (2019).

23. Makowski, E. K. et al. Optimization of therapeutic antibodies for reduced self-association and non-specific binding via interpretable machine learning. *Nat*. Biomed. Eng. 8, 45–56 (2024).

24. Lippow, S. M., Wittrup, K. D. & Tidor, B. Computational design of antibody-affinity improvement beyond in vivo maturation. Nat. Biotechnol. 25, 1171–1176 (2007).

25. Warszawski, S. et al. Optimizing antibody affinity and stability by the automated design of the variable light-heavy chain interfaces. PLoS Comput. Biol. 15, e1007207 (2019).

26. Borenstein-Katz, A. et al. Biomolecular recognition of the glycan neoantigen CA19-9 by distinct antibodies. J. Mol. Biol. 433, 167099 (2021).

27. Rosace, A. et al. Automated optimisation of solubility and conformational stability of antibodies and proteins. Nat. Commun. 14, 1937 (2023).

28. Tennenhouse, A. et al. Energy-guided combinatorial co-optimization of antibody affinity and stability. bioRxiv (2025) doi:10.1101/2025.11.26.690765.

29. Bennett, N. R. et al. Atomically accurate de novo design of antibodies with RFdiffusion. Nature 649, 183–193 (2026).

30. Mille-Fragoso, L. S. et al. Efficient generation of epitope-targeted antibodies with Germinal. Nat. Biotechnol. (2026) doi:10.1038/s41587-026-03187-0.

31. Balbi, P. E. M. et al. Mapping targetable sites on the human surfaceome for the design of novel binders. Proc. Natl. Acad. Sci. U. S. A. 123, e2506269123 (2026).

32. Zhou, J., Le, C. Q., Zhang, Y. & Wells, J. A. A general approach for selection of epitope-directed binders to proteins. Proc. Natl. Acad. Sci. U. S. A. 121, e2317307121 (2024).

33. Shrock, E. L. et al. Germline-encoded amino acid-binding motifs drive immunodominant public antibody responses. Science 380, eadc9498 (2023).

34. Lipsh-Sokolik, R. et al. Combinatorial assembly and design of enzymes. Science 379, 195–201 (2023).

35. Tsuboyama, K. et al. Mega-scale experimental analysis of protein folding stability in biology and design. Nature 620, 434–444 (2023).

36. Weinstein, J. Y. et al. Designed active-site library reveals thousands of functional GFP variants. Nat. Commun. 14, 2890 (2023).

37. Dunbar, J. et al. SAbDab: the structural antibody database. Nucleic Acids Res. 42, D1140–6 (2014).

38. Abanades, B. et al. ImmuneBuilder: Deep-Learning models for predicting the structures of immune proteins. *Commun*. Biol. 6, 575 (2023).

39. Engler, C., Kandzia, R. & Marillonnet, S. A one pot, one step, precision cloning method with high throughput capability. PLoS One 3, e3647 (2008).

40. Ariel. CADAbRe. Preprint at 10.17605/OSF.IO/CAZV7 (2025).

41. Alford, R. F. et al. The Rosetta all-atom energy function for macromolecular modeling and design. J. Chem. Theory Comput. 13, 3031–3048 (2017).

42. Zemlin, M. et al. Expressed murine and human CDR-H3 intervals of equal length exhibit distinct repertoires that differ in their amino acid composition and predicted range of structures. J. Mol. Biol. 334, 733–749 (2003).

43. Maciá Valero, A., Prins, R. C., de Vroet, T. & Billerbeck, S. Combining oligo pools and Golden Gate cloning to create protein variant libraries or guide RNA libraries for CRISPR applications. Methods Mol. Biol. 2850, 265–295 (2025).

44. Andris-Widhopf, J., Rader, C., Steinberger, P., Fuller, R. & Barbas, C. F., 3rd. Methods for the generation of chicken monoclonal antibody fragments by phage display. J. Immunol. Methods 242, 159–181 (2000).

45. Tsapogas, P., Mooney, C. J., Brown, G. & Rolink, A. The cytokine Flt3-ligand in normal and malignant hematopoiesis. Int. J. Mol. Sci. 18, E1115 (2017).

46. Wilson, K. R., Villadangos, J. A. & Mintern, J. D. Dendritic cell Flt3 - regulation, roles and repercussions for immunotherapy. Immunol. Cell Biol. 99, 962–971 (2021).

47. Madsen, A. V., Pedersen, L. E., Kristensen, P. & Goletz, S. Design and engineering of bispecific antibodies: insights and practical considerations. Front. Bioeng. Biotechnol. 12, 1352014 (2024).

48. Ferrara, F. et al. A pandemic-enabled comparison of discovery platforms demonstrates a naïve antibody library can match the best immune-sourced antibodies. Nat. Commun. 13, 462 (2022).

49. Cauerhff, A., Goldbaum, F. A. & Braden, B. C. Structural mechanism for affinity maturation of an anti-lysozyme antibody. Proc. Natl. Acad. Sci. U. S. A. 101, 3539–3544 (2004).

50. Jain, T. et al. Assessment and incorporation of in vitro correlates to pharmacokinetic outcomes in antibody developability workflows. MAbs 16, 2384104 (2024).

51. Jain, T. et al. Assessment and incorporation of in vitro correlates to pharmacokinetic outcomes in antibody developability workflows. MAbs 16, 2384104 (2024).

52. Hötzel, I. et al. A strategy for risk mitigation of antibodies with fast clearance. MAbs 4, 753–760 (2012).

53. Jacobs, S. A., Wu, S.-J., Feng, Y., Bethea, D. & O’Neil, K. T. Cross-interaction chromatography: a rapid method to identify highly soluble monoclonal antibody candidates. Pharm. Res. 27, 65–71 (2010).

54. Geng, S. B., Cheung, J. K., Narasimhan, C., Shameem, M. & Tessier, P. M. Improving monoclonal antibody selection and engineering using measurements of colloidal protein interactions. J. Pharm. Sci. 103, 3356–3363 (2014).

55. Liu, Y. et al. High-throughput screening for developability during early-stage antibody discovery using self-interaction nanoparticle spectroscopy. MAbs 6, 483–492 (2014).

56. Sule, S. V. et al. High-throughput analysis of concentration-dependent antibody self-association. Biophys. J. 101, 1749–1757 (2011).

57. DeLaitsch, A. T. et al. Selective recognition of carbohydrate antigens by germline antibodies isolated from AID knockout mice. J. Am. Chem. Soc. 144, 4925–4941 (2022).

58. Griffiths, A. D. et al. Isolation of high affinity human antibodies directly from large synthetic repertoires. EMBO J. 13, 3245–3260 (1994).

59. Nissim, A. et al. Antibody fragments from a ‘single pot’ phage display library as immunochemical reagents. EMBO J. 13, 692–698 (1994).

60. Pliakostathis et al. EUR 30185 - EURL ECVAM Recommendation on Non-Animal Derived Antibodies. Preprint at 10.2867/28061.

61. Bradbury, A. & Plückthun, A. Reproducibility: Standardize antibodies used in research. Nature 518, 27–29 (2015).

62. Wilson, I. A. & Stanfield, R. L. 50 Years of structural immunology. J. Biol. Chem. 296, 100745 (2021).

63. Sheriff, S. et al. Three-dimensional structure of an antibody-antigen complex. Proc. Natl. Acad. Sci. U. S. A. 84, 8075–8079 (1987).

64. Abhinandan, K. R. & Martin, A. C. R. Analysis and improvements to Kabat and structurally correct numbering of antibody variable domains. Mol. Immunol. 45, 3832–3839 (2008).

65. Kabat, E. A. & Wu, T. T. Attempts to locate complementarity-determining residues in the variable positions of light and heavy chains. Ann. N. Y. Acad. Sci. 190, 382–393 (1971).

66. Fleishman, S. J. et al. RosettaScripts: a scripting language interface to the Rosetta macromolecular modeling suite. PLoS One 6, e20161 (2011).

67. Giudicelli, V., Chaume, D. & Lefranc, M.-P. IMGT/GENE-DB: a comprehensive database for human and mouse immunoglobulin and T cell receptor genes. Nucleic Acids Res. 33, D256–61 (2005).

68. Oomen, C. J. et al. Crystal structure of an Anti-meningococcal subtype P1.4 PorA antibody provides basis for peptide-vaccine design. J. Mol. Biol. 351, 1070–1080 (2005).

69. Lipsh-Sokolik, R., Listov, D. & Fleishman, S. J. The AbDesign computational pipeline for modular backbone assembly and design of binders and enzymes. Protein Sci. 30, 151–159 (2021).

70. Raybould, M. I. J. et al. Five computational developability guidelines for therapeutic antibody profiling. Proc. Natl. Acad. Sci. U. S. A. 116, 4025–4030 (2019).

71. Shirai, H., Kidera, A. & Nakamura, H. H3-rules: identification of CDR-H3 structures in antibodies. FEBS Lett. 455, 188–197 (1999).

72. Zulkower, V. & Rosser, S. DNA Chisel, a versatile sequence optimizer. Bioinformatics 36, 4508–4509 (2020).

73. Harris, C. R. et al. Array programming with NumPy. Nature 585, 357–362 (2020).

74. Subramanian, S. K., Russ, W. P. & Ranganathan, R. A set of experimentally validated, mutually orthogonal primers for combinatorially specifying genetic components. Synth. Biol. 3, (2018).

75. Cock, P. J. A. et al. Biopython: freely available Python tools for computational molecular biology and bioinformatics. Bioinformatics 25, 1422–1423 (2009).

76. Helsmoortel, M. et al. Oxford Nanopore enhanced accuracy of long-read amplicons applied to microbial whole-genome sequencing. Microbiol. Spectr. 14, e0285625 (2026).

77. Li, H. et al. The Sequence Alignment/Map format and SAMtools. Bioinformatics 25, 2078–2079 (2009).

78. Li, H. Minimap2: pairwise alignment for nucleotide sequences. Bioinformatics 34, 3094–3100 (2018).

79. Ewing, B., Hillier, L., Wendl, M. C. & Green, P. Base-calling of automated sequencer traces using phred. I. Accuracy assessment. Genome Res. 8, 175–185 (1998).

80. Altschul, S. F., Gish, W., Miller, W., Myers, E. W. & Lipman, D. J. Basic local alignment search tool. J. Mol. Biol. 215, 403–410 (1990).

81. Baker, N. A., Sept, D., Joseph, S., Holst, M. J. & McCammon, J. A. Electrostatics of nanosystems: application to microtubules and the ribosome. Proc. Natl. Acad. Sci. U. S. A. 98, 10037–10041 (2001).

82. Slezak, T. et al. Engineered protein G variants for multifunctional antibody-based assemblies. Protein Sci. 34, e70019 (2025).

83. Gasteiger, E. et al. Protein identification and analysis tools on the ExPASy server. in The Proteomics Protocols Handbook 571–607 (Humana Press, Totowa, NJ, 2005).

84. Abramson, J. et al. Accurate structure prediction of biomolecular interactions with AlphaFold 3. Nature 630, 493–500 (2024).

85. Meng, E. C. et al. UCSF ChimeraX: Tools for structure building and analysis. Protein Sci. 32, e4792 (2023).

86. Emsley, P., Lohkamp, B., Scott, W. G. & Cowtan, K. Features and development of coot. *Acta Crystallogr*. D Biol. Crystallogr. 66, 486–501 (2010).

87. Adams, P. D. et al. PHENIX: a comprehensive Python-based system for macromolecular structure solution. Acta Crystallogr. D Biol. Crystallogr. 66, 213–221 (2010).

